# Duckweed evolution: from land back to water

**DOI:** 10.1101/2023.03.22.533731

**Authors:** Yang Fang, Xueping Tian, Yanling Jin, Anping Du, Yanqiang Ding, Zhihua Liao, Kaize He, Yonggui Zhao, Ling Guo, Yao Xiao, Yaliang Xu, Shuang Chen, Yuqing Che, Li Tan, Songhu Wang, Jiatang Li, Zhuolin Yi, Lanchai Chen, Leyi Zhao, Fangyuan Zhang, Guoyou Li, Jinmeng Li, Qinli Xiong, Yongmei Zhang, Qing Zhang, Xuan Hieu Cao, Hai Zhao

## Abstract

Terrestialization is supposedly an important evolutionary process plant experience. However, directions of land back to water acquired little attention. Here we integrate multiproxy evidence to elucidate the evolution of duckweed. Three genera of duckweed show chronologically gradient degeneration in roots structure and stomatal function and decrease in lignocellulose content, accompanied by gradual contraction in relevant gene numbers and/or decline in transcription. The gene numbers in the main phytohormonal pathway are also gradually decreased. The co-action of auxin and rhizoid development gene causes a gradual decrease in adventitious roots. The significant expansion of the flavonoid pathway is also related to the adaptation of duckweed to floating growth. This study reconstructs the evolution history of duckweeds from land back to water, reverse to that of early land plants.

**Summary:** With terrestrialization being the popularly acknowledged plant evolutionary process, little is known about the evolution of higher plant from land back to water. Here we integrate multiproxy evidence to elucidate the gradual reverse evolution of duckweed. Three genera of duckweed show chronologically gradient degeneration in the structure of roots, the function of stomata, and decrease in lignocellulose content, accompanied by gradient contraction in relevant gene numbers and/or decline in transcript expression. The gene numbers in the main phytohormonal pathway are also gradually decreased. The co-action of auxin and rhizoid development gene causes a gradual decrease in adventitious roots. The significant expansion of the flavonoid pathway is also highly related to the adaptation of duckweed to floating growth. Our study combined with the fossil evidence reconstruct the evolution history of duckweeds from land back to water, reverse to that of early land plants. This study reconstructed the process of how a land plant returns to water, a reverse evolutionary approach which is different from what we studied in textbook about plant terrestrialization. This finding could be helpful for us to deeply and widely understand the adaptation of plant to the environment, and to expand and deepen the knowledge of evolution theory.

## 1 Introduction

Life on Earth originated in water 4.0-3.8 billion years ago, where aquatic algae first transformed from water to land 450 million years ago (MYA)^1^. Terrestrialization was first initiated by plants and then followed by animals. During the process, four main traits in plants concerning land adaptability started to emerge around 400 MYA, including stomata^2^, lignin^3^, roots^4^ and more phytohormones^5^. The occurrence of such new characteristics brought about tremendous changes in plant life forms and expanded biodiversity on Earth. In fact, the number of species and total biomass of terrestrial organisms far exceeded that of aquatic ones. Although land only accounts for 29% of the total Earth’s surface, just terrestrial angiosperms contains around 400,000 species, equivalent to more than 4 times of the number of aquatic plants^6^. Meanwhile, the biomass of land plants accounts for 82% of the total biosphere biomass^7^.

These evolutionary phenomenon from water to land has always intrigued scientists in relative area^5^. However, the reversed evolution from land back into water has not been studied much. Even though there were studies confirming such process, most of the evidence derived from only molecular phylogenetic trees established with some genes of aquatic plants^8–10^. These studies lacked evidence on how the main characteristics, functions, and genes of these plants adapted to water.

Duckweed may be the answer to reversed terrestrialization. The floating monocot and smallest flowering plant on Earth comprises of two subfamilies, Lemnoideae and Wolffioideae, which further divide into three and two genera respectively, *Landoltia*, *Lemna*, *Spirodela* and *Wolffia* and *Wolffiella*^11^. Evolutionarily, among the five genera, *Spirodela* was the first to appear in duckweed subfamily, followed by *Landoltia*, *Lemna*, *Wolffiella*, and *Wolffia* subsequently. Surprisingly, when examining them in sequence, we discovered trends of diminishing organism size and disappearing root number. This suggests the possibility of reversed terristrialization. Here, we selected *Spirodela polyrhiza* strain 7498, *Landoltia punctata* strain 0202, and *Lemna minor* strain 7753 from their respective genera and integrated their omics, morphological and physiological data. Our results showed that the four most important terrestrial traits, roots, stomata, hormone, and lignocellulose, are degrading or even vanishing in aspects of genome, transcription, function, structure, and composition.

This study extant and fossil (*Limnobiophyllum scutatum*)^12^ data reconstruct the evolution of duckweed from land back to water and achieves such result using extant species.

## 2 Results and discussion

### 2.1 Duckweed phylogeny

We sequenced, assembled, and annotated the *L. punctata* genome (GenBank assembly accession: PRJNA546087) and did comparative genomic analysis with other two published duckweed genomes, *S. polyrhiza* and *L. minor*, as well as genomes of some model species and evolutionary node species including *Klebsormidium flaccidum*, *Zostera marina*, *Arabidopsis thaliana*, *Oryza sativa* and *Zea mays* (Supplementary Table 11). A phylogenetic tree was constructed using genes of 1545 single-copy families (Supplementary Data 19) shared by these eight species to determine their evolutionary relationship with duckweed. Our results were consistent with previous studies on phylogeny and systematics of duckweed subfamily^13, 14^. Meanwhile, the comparative transcriptome was also analyzed among *S. polyrhiza*, *L. punctata,* and *L. minor* to further understand their function and structure.

Compared with the algae *K. flaccidum* that contains some land-related genes, land plants (*Z. mays, A. thaliana, O. sativa*) exhibits a higher number of genes relevant to terrestrial traits including root, stomata, and hormone. On the contrary, the total numbers of genes related to these traits in aquatic duckweeds (*S. polyrhiza*, *L. punctata*, *L. minor*) significantly reduced compared to typical land plants (Fig. 1b). As a kind of angiosperm that belongs to Araceae, Alismatales, duckweed had early divergences with the basal lineage of Alismatales (*Tofieldia thibetica*) at approximately 120.0 MYA, and had early divergences with basal lineage of Araceae (*Symplocarpus renifolius*) at approximately 110.0 MYA^15, 16^, and then became diverse at approximately 63.4 MYA and further differentiated at 46.7 MYA (Fig. 1b, Extended Data Fig.1). Furthermore, those genes related to terrestrial characteristics of roots, stomata, and hormones gradually reduced along the evolutionary sequence of duckweeds (Fig. 1b).

**Fig. 1.**
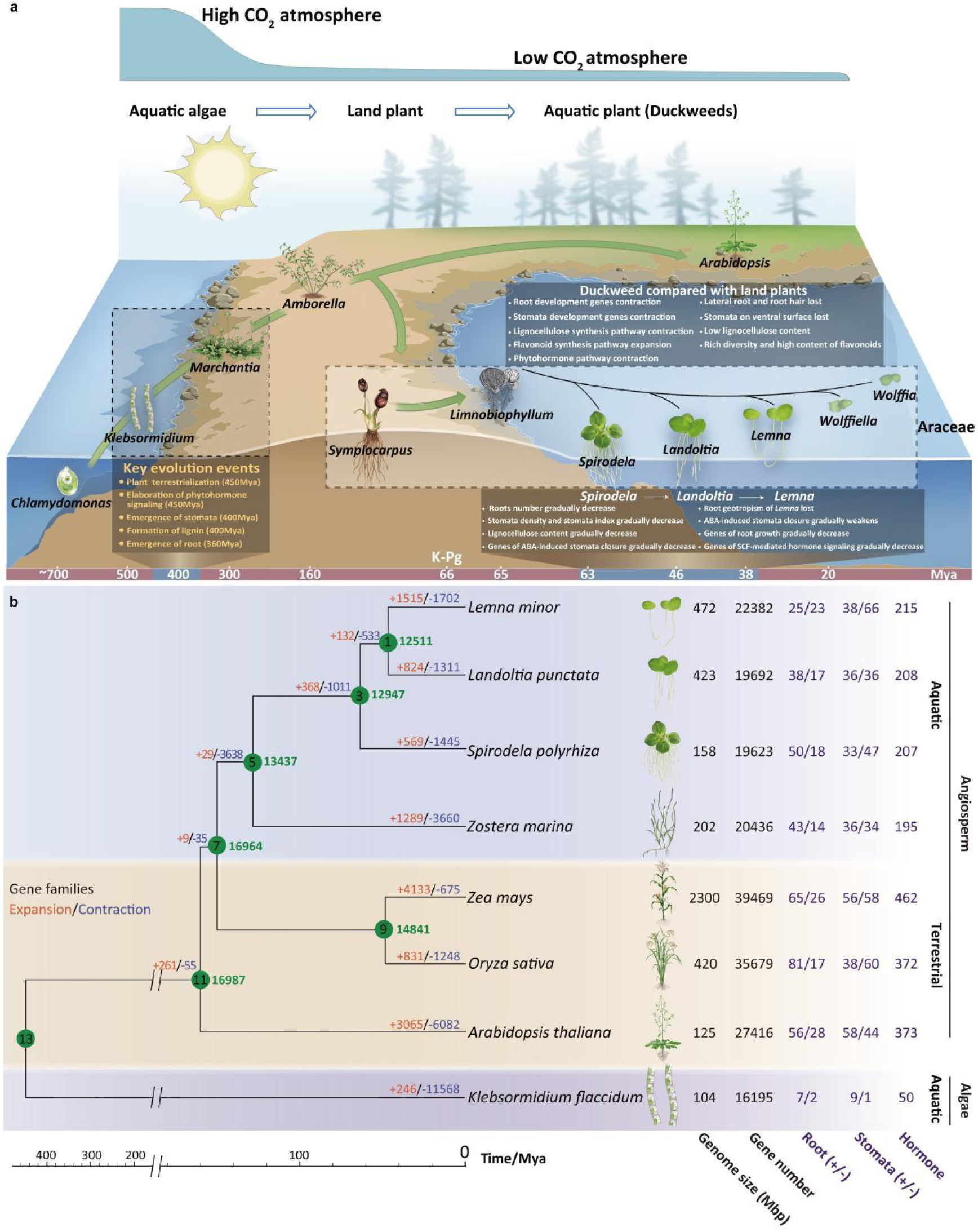
The evolution model, phylogenetic tree and gene family analysis of duckweeds. **a**, Evolution of duckweed from land back to water. **b,** Phylogenetic tree of duckweeds and its gene family analysis compared with 5 representatives of the Viridiplantae. Phylogenetic context of duckweeds and the number of gene copies involved in root development (Root), stomata development (Stomata), and phytohormone pathways (Hormone). +, positively related to development; -, negatively related to development. The numbers at the nodes of the phylogenetic tree indicate the divergence time in MYA.

Cretaceous-Paleogene (K-Pg) extinction (66.0 MYA) is one of the most important events in the evolution of life on Earth. It provided empty ecological niches and induced adaptive radiations to numerous organisms^17–19^. For example, the extinction of dinosaurs and the emergence of birds are closely related to the evolution of adaptive radiations^20, 21^. Interestingly, duckweed first differentiated into *Spirodela* (63.4 MYA) after K-Pg extinction, followed by *Landoltia* and *Lemna* genus, and finally *Wolffiella* and *Wolffia* (Fig. 1a). The evolution of duckweed is also possibly driven by adaptive radiation after K-Pg extinction.

### 2.2 Root traits and development

#### 2..2.1 Disappearing roots in duckweeds

The emergence of roots during the Devonian Period (416-360 MYA) was essential for plants to colonize the land^22^. Sophisticated root system^23^ in land plants is to anchor into soil, sense gravity, acquire nutrients and water^4^. Duckweed belongs to the Araceae family, whose terrestrial basal lineage *Symplocarpus*^16^ (Extended Data Fig.1) has large root systems^24^. Interestingly, the fossil of duckweeds’ ancestor *L. scutatum* indicated that they were also floating aquatic angiosperm with structure like stolon, numerous adventitious roots^25–30^, some bore stout primary roots (3 mm wide), and lateral roots^12, 31^. Such characteristics exhibits a transitional state between terrestrial ancestor (Supplementary Fig. 7) and extant duckweeds. The root traits of extant duckweeds are highly reduced compared with the terrestrial basal group of Araceae and their fossil ancestor. Some studies proved weakened root absorption in duckweed and suggested anchoring as the main function^32^. The numbers of genes related to root development in duckweeds (Supplementary Data 1) not only remarkably reduced compared with *A. thaliana*, *O. Sativa* and *Z. mays*, but also gradually contracted by the evolutionary sequence of the three species of duckweed (Fig. 1b). *S. polyrhiza*, *L. punctata,* and *L. minor* have 50, 38, and 25 genes of positively regulated root development, and 18, 17, and 23 genes of negatively regulated root development (Fig. 1b). Moreover, the expressions of these positive genes are generally higher in *S. polyrhiza* than those in *L. punctata* and *L. minor* (Extended Data Fig.2).

**Fig. 2.**
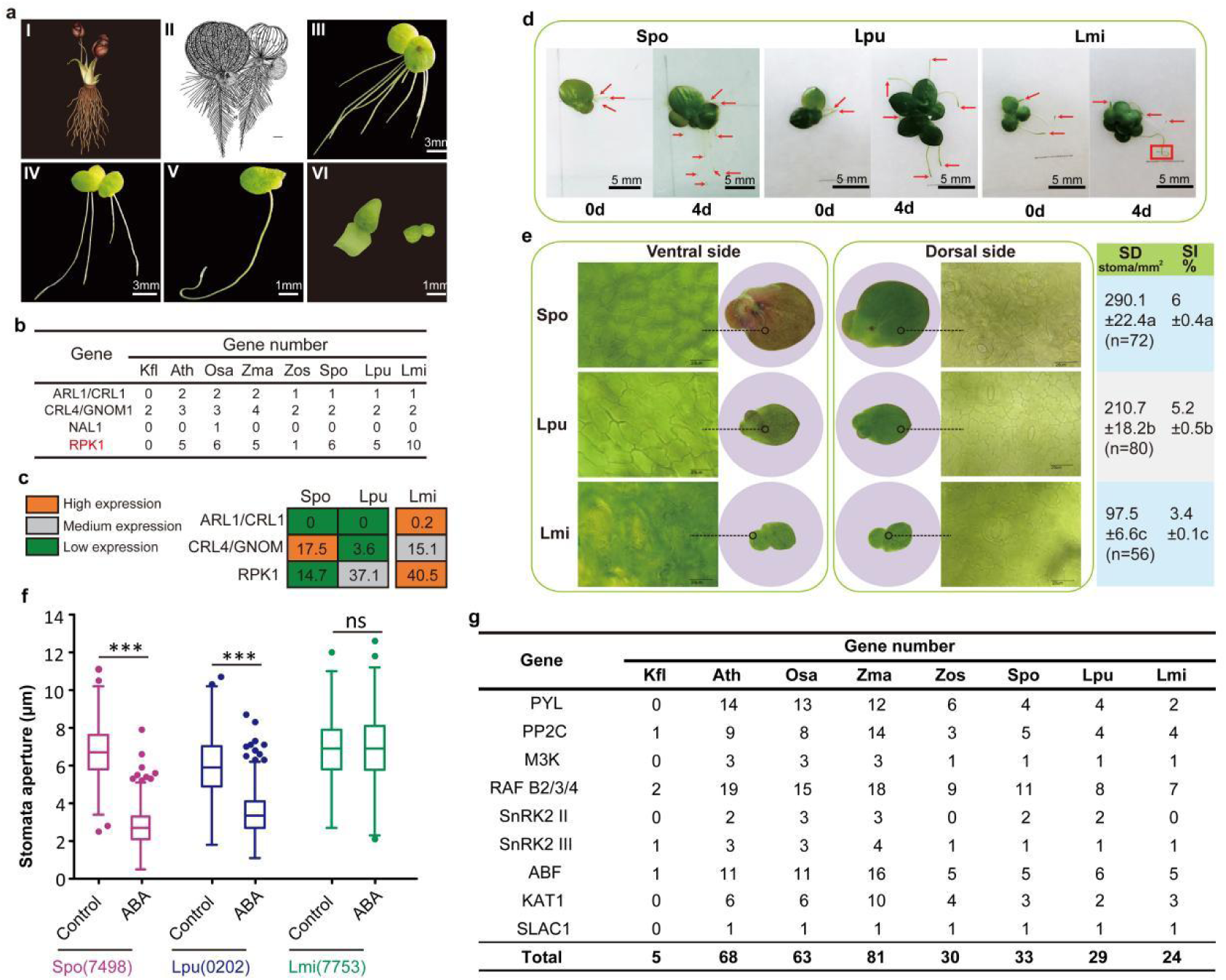
Genetic development and physiological characteristics of root and stomata of duckweeds. **a**, Root numbers and morphology of *Symplocarpus* (I), the fossil of *Limnobiophyllum scutatum* (ancestors of duckweed family)^12^ (II) and duckweeds (III-VI). Root numbers: *S. polyrhiza* >7 (III), *L. punctata* 2-7 (IV), *L. minor* 1 (V), *Wolffiella* genus (left of VI) and *Wolffia* genus (right of VI) have no root. **b**, The number of key genes in adventitious root emergence and development of duckweeds and other plants. Red indicates negative regulation. **d,** The geotropism of *S. polyrhiza* (Spo), *L. punctata* (Lpu), and *L. minor* (Lmi) roots in different culture time. The red arrows indicate duckweed roots. The direction of gravity is vertical downward. **c**, The expression of key genes in adventitious root emergence and development of duckweeds. **e**, Stomata distribution on dorsal and ventral sides of duckweed fronds. Stomata were monitored on the whole duckweed fronds. SD, stomatal density, SD (stomata/*mm^2^* = 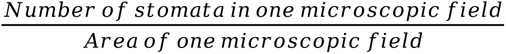 SI, stomatal index, SI % = 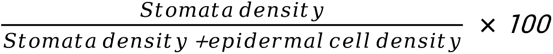; *n*, the total number of surveyed stomata. Values represent mean ± standard deviations. Letters indicate significant differences determined by one-way ANOVA followed by multiple comparison with Tukey-Kramer test (p<0.05). **f,** Comparison of ABA-induced stomatal closure in *S. polyrhiza* (Spo 7498), *L. punctata* (Lpu 0202), and *L. minor* (Lmi 7753). The duckweeds were placed in 1/5 Hoagland medium for 3 h under white light to induce stomata opening and further incubated in the absence (Control) or in the presence 100 μM ABA (ABA). Stomatal aperture was measured after incubating for 60 min in room temperature when 100 μM ABA was added. More than 400 stomata were measured for each species. Y-axis represents mean ± standard deviations; Bars = 10 μm. ***, P<0.001; ns, not significant. **g,** The number of genes involved in ABA-induced stomatal closure signaling in eight typical species. Kfl, *K. flaccidum*; Ath, *A. thaliana*; Osa, *O. Sativa*; Zma, *Z. mays*; Zos, *Z. marina*; Spo, *S. polyrhiza*; Lpu, *L. punctata*; Lmi, *L. minor*.

Extant duckweed only has adventitious root. Numbers of the roots on different duckweed genera decrease as follows: over 7 roots in *Spirodela*, 2-7 roots in *Landoltia*, only 1 root in *Lemna*, and no root in *Wolffiella* and *Wolffia* (Fig. 2a)^33^. The number of key genes of adventitious root emergence and development is contracted (*ARL1*/*CRL1*, *CRL4*/*GNOM1*) or lost (*NAL1*) in duckweeds. The negative regulatory gene *RPK1* was not less than *A. thaliana*, *O. sativa,* and *Z. mays*, and the number was even expanded in *Lemna*. The *ARL1*/*CRL1* genes was almost not expressed in duckweeds, and *CRL4*/*GNOM1* in *Spirodela* was higher than that in *Landoltia* and *Lemna.* Meanwhile, *RPK1* gene in *Lemna* showed the highest expression level amongst the three species (Fig.2b, c, Supplementary Data 16). Therefore, the numbers of key genes and their expression levels involved in adventitious root development gradually decrease along the evolution direction of duckweeds, confirming the trend of diminishing adventitious root in duckweed.

Auxin response in land plants largely determines their morphological traits adapting to terrestrial habitats, particularly rhizoid development and organized three-dimensional growth. The emergence and expansion of the auxin response system aligns well with the morphological changes controlled by auxin^34^. However, the highly degenerated and almost two-dimensional structure in duckweed confirm the contraction of their auxin response system. Auxin promotes the formation and development of adventitious root through hormone signaling. Along the evolution direction of duckweed subfamily, the number and expression level of key gene *T1R1/AFB* decreases while those of the negative regulatory gene *RPK1* increases. The number of *PIN* in *L. minor* (8) is higher than that in *S. polyrhiza* (4) and *L. punctata* (4), and their expression levels remarkably decrease along the evolution direction (FPKM=80.4, 17.9, 2.1) (Supplementary Data 16). This also indicated that the cross-talking between auxin and root development genes leads to a gradient decrease in the number of roots and a gradual weakening root function in the process of migrating back to water (Fig. 3b).

**Fig. 3.**
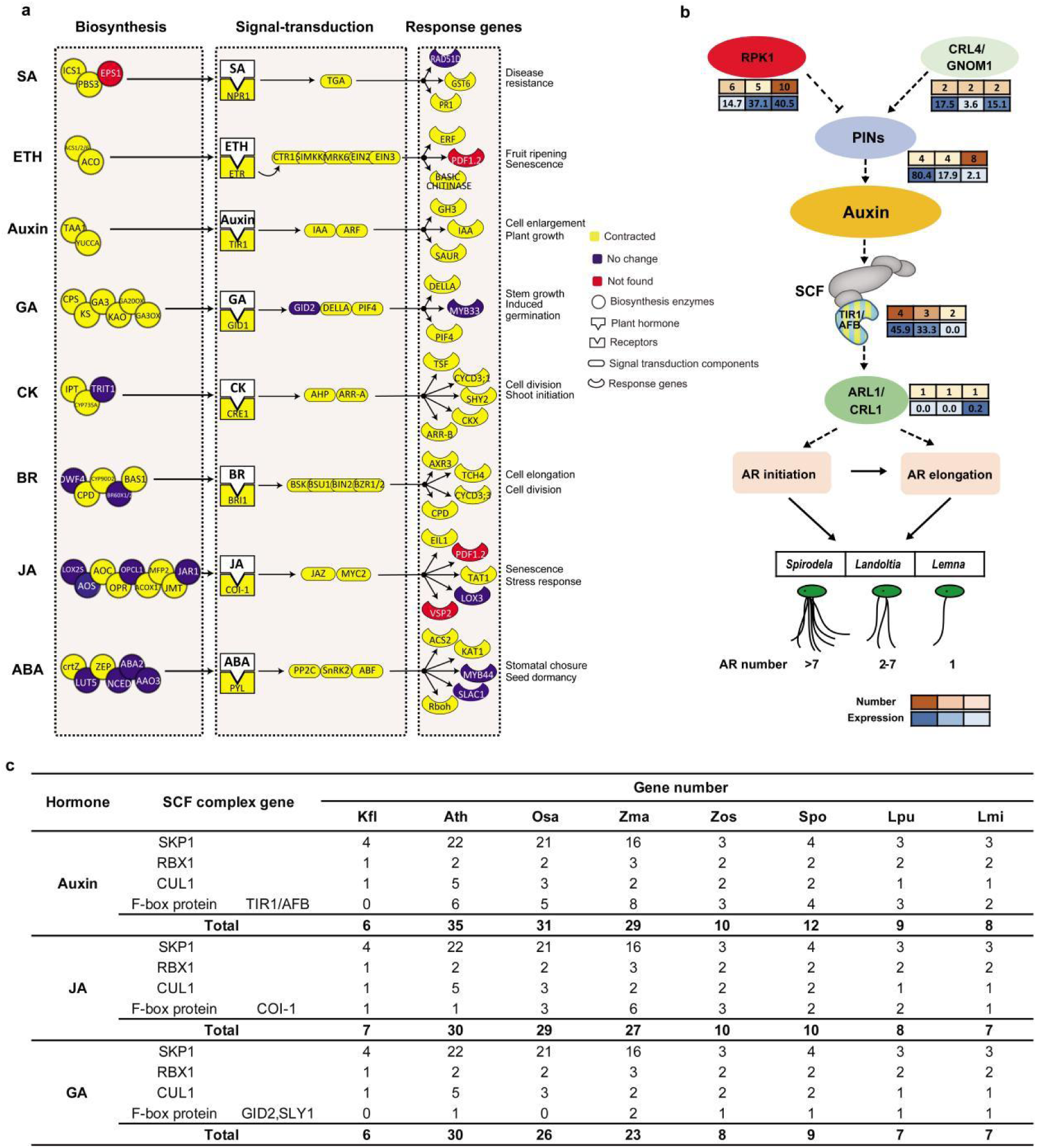
Phytohormones pathways and the cross talking between auxin and AR development in duckweeds. **a**, Comparison of phytohormones pathways between duckweeds and typical land plants. Mean numbers of genes in *S. polyrhiza*, *L. punctata* and *L. minor* were compared to those of *A. thaliana*, *O. sativa*, and *Z. mays*. The number of genes obtained from gene family analysis via OrthMCL. The compared genes involve in biosynthesis, signal transduction, and response of phytohormones. Detailed information about the gene number and gene ID is provided in Supplementary Data 3. **b,** The combined effects of auxin and rhizoid development gene on adventitious roots of duckweeds. The heatmaps next to genes represent the corresponding gene number and its expression (FPKM value). The upper panels represent gene number, and the lower panels represent corresponding gene expression. In SCF complex, only the number and expression of TIR1/AFB genes are shown. Arrows and dotted arrows represent positive regulatory actions, and dotted lines ending in flat heads indicate negative regulatory actions. SCF, Skp1/Cullin/F-box-type ubiquitin ligase; AR, adventitious root. **c,** The number of genes involved in Skp1/Cullin/F-box (SCF)-mediated hormone signaling of auxin, jasmonic acid (JA) and gibberellic acid (GA).

#### 2.2.2 Lateral roots and root hairs in duckweeds

Lateral roots and root hairs are not found in extant duckweeds, but fossil records of *L. scutatum,* the ancestor of duckweed, proved their previous existance^12, 31^. The genes or transcription factors (TFs) related to the formation of lateral roots and root hairs are lost (*LBD16*, *CPC* and *WER*) or overall contracted (*WAK*, *LBD29*, *GL3* and *GL2*) (Supplementary Data 1) in the three species. Previous studies on *S. polyrhiza* and rice focused on the genes of phytohormone signal transduction in lateral roots and root hair elongation^32, 35^, whereas this study specifically analyzed the loss and contraction of key genes involved in lateral root formation and epidermal cell differentiation of root hair in the three species.

#### 2.2.3 Lost gravity response of *Lemna*

Generally, roots of land plants grow towards gravity to anchor into soil^36^, however, the roots of *L. minor* grow upward (Fig. 2d). Although the three species of duckweeds possess three regulatory genes (*PLC1*, *WAV3*, and *AGR*) important for gravitational responses, the number of *PLC1* is reduced from 4-7 in model plants to 2 (Supplementary Data 1). More importantly, expression of these genes were not detected in *L. minor* (Extended Data Fig.2), which may be the reason for the loss of its gravity response (Fig. 2d).

#### 2.2.4 Stomata distribution and development

Plant stomata facilitates adaptation to terrestrial environment for its function of optimizing gas exchange and limiting evaporative loss^2, 25^. The earliest land plants lacked stomata and were restricted to moist habitats. Around 400 MYA, stomata appeared firstly on shade-tolerant terrestrial species such as vascular plants. Right after, plants flourished on land^28^. Therefore, stomata is an important adaptive trait for transition to land.

Further, stomata density (SD) and stomata index (SI) of plants are indicators of the level of adaptation to environment^30, 37^. Duckweed stomata exist only on the dorsal surface (Fig. 2e), differing from most land plant leaves that includes stomata on both the ventral and dorsal^29^. In its transition out of water, duckweed’s stomata also experienced adaptive evolution. *S. polyrhiza*, *L. punctata*, and *L. minor* reach SD 290.1, 210.7, and 97.5 stomata mm^-2^, and achieve SI 6.0 %, 5.2 %, and 3.4 %, respectively (Fig. 2e). Both SD and SI of the three species exhibit similar declining (Fig. 1b). The numbers of genes involved with stomatal development in these three species exhibits similar trend to their SD and SI (Supplementary Data 2). The number of TF *SCRM*, one of the most important genes involved in stomata lineage progression, decreases to one copy in *L. punctata* and *L. minor*, and two copies in *S. polyrhiza*. Likewise, EPFL9 (positive regulation of stomatal complex development) and *CDKB*1;1 (guard mother cell differentiation) decreased to one copy in *S. polyrhiza* and *L. punctata* and no copies in *L. minor*(Supplementary Data 2). Compared to *S. polyrhiza* and *L. punctata*, *L. minor* has much more genes suppressing stomal generation with higher expression levels in some, such as *SDD*1 and *MPK3/MPK6* (Extended Data Fig.3). These genomic and transcriptome results are consistent with the morphological changes of stomata within the duckweed subfamily.

### 2.3 Phytohormone pathways

#### 2.3.1 Phytohormone pathways contraction

Terrestrial environments are more complicated and varying than aquatic ones. Phytohormones in plants play a primary role in adapting to the complex land environment^38^. Although aquatic algae already possessed complete biosynthesis pathways of major phytohormones before migrating to land^39^, their signaling transduction pathways remained incomplete. The earliest land plants such as charophytes and liverworts (*K. flaccidum*, *Chara braunii*, and *Marchantia polymorpha*) first expanded these biosynthesis pathways and started to complete the signaling transduction pathways^5, 40, 41^. Both processes were concluded with the emergence of angiosperm. Duckweeds possess the complete biosynthesis pathway, signal transduction pathway, and response genes of eight major phytohormones including ABA, SA, JA, ethylene (ETH), auxin (Aux), gibberellic acid (GA), cytokinine (CK), Brassinosteroid (BR) (Fig. 3a, Supplementary Fig. 8). All listed phytohormones were detectable (Supplementary Fig. 8). However, compared with that in land angiosperms such as *A. thaliana*, *O. sativa*, and *Z. mays*, the gene number of signaling transduction pathway and response gene number are remarkably lower in duckweed (Fig. 3a). A sharp contrast is seen in this contraction and the expansion of the phytohormone pathway in the earliest land plants.

#### 2.3.2 Gradient reduction of SCF-Mediated Hormone Signaling

Auxin, JA, and GA are a group of phytohormones. Their most important common feature is that proteolytic targeting is mediated by Skp1 / Cullin / F-box (SCF) E3 ubiquitin ligase complex. SCF complex of auxin, JA and GA appears first in the land plant lineage^34^. In algae *K. flaccidum* that first acquired the fundamental mechanism to adapt to terrestrial environments, only JA had a complete SCF complex genes, both auxin and GA lacked the gene of F-box protein. Meanwhile, *K. flaccidum* has only 6, 7, and 6 SCF complex genes of auxin, JA, and GA signaling pathways, respectively (Fig. 3c). Then the gene numbers of SCF complex gradually increased along the terrestrialization of plants, especially that of the auxin response system. Duckweeds have complete SCF complex genes of auxin, JA, and GA, but the gene numbers contract significantly compared to land plants. The gene numbers of auxin SCF complex are 12, 9 and 8 in *S. polyrhiza*, *L. punctata* and *L. minor*, respectively. Their numbers remarkably reduced compared with *A. thaliana* (35), *O. Sativa* (31) and *Z. mays* (29) and gradually contracted along the three species in the listed order (Fig. 3c). Moreover, the gene numbers of SCF complex of auxin, JA and GA in *S. polyrhiza*, *L. punctata,* and *L. minor* are also remarkably reduced compared with *A. thaliana*, *O. Sativa,* and *Z. mays*, and also gradually contracted in these three species of duckweed (Supplementary Data 14).

#### 2.3.3 Gradually weakened function of ABA-induced stomatal closure

The regulation of ABA in response to drought stress is the most important function of land plants and is mainly achieved by adjusting the closure of stomata^42^. Stomata and its ABA-induced closure emerged in early land plants such as mosses and lycophytes and further developed in ferns^43^. The stomata of typical angiosperms can close completely under ABA treatment. ABA-induced stomatal closure exists in duckweeds but weakens gradually along the evolution direction of the three species. Surprisingly, unlike other typical angiosperms, the stomata of *S. polyrhiza* and *L. punctata* cannot close completely in response to ABA. This trait is also observed in an early land moss (*Physcomitrella patens*)^44^ (Fig. 2f). More importantly, the level of stomatal closure from ABA treatment decreases in the order of *S. polyrhiza* (stomata aperture from 6.7±1.4 μm to 2.8±1.0 μm), *L. punctata* (stomata aperture from 6.0 ± 1.6 μm to 3.5 ± 1.2 μm), *L. minor* (stomata aperture from 6.9 ±1.4μm to 6.9 ±1.8μm). In fact, the last species was completely unaffected by ABA treatment (Fig. 2f).

K. *flaccidum* has no stomata and no complete genes (only 5 genes) regulating ABA-induced stomatal closure signaling pathway. Evolutionarily, these gene peaked with the emergence of angiosperm, reaching 68, 63, and 81 genes in *A. thaliana*, *O. sativa,* and *Z. mays* (Fig. 2g). Compared with land plants, the gene numbers of this pathway in all three duckweed species are contracted and follows their evolution direction (33, 29, 24 respectively) (Fig. 2g, Supplementary Data 15). This indicated that in duckweeds, the decreasing trend of gene numbers in ABA-induced stomatal closure signaling was consistent with the level of their closure in the three species.

In the three species of duckweeds, the number of genes significantly contracted in major phytohormone pathways, contrary to the expansion trend in the land plants. In particular, along the evolutionary sequence of duckweed, not only did the stomatal development of the three species gradually weaken, but the response of stomata to ABA treatment also gradually weakened from *S. polyrhiza* to *L. punctata* and to *L. minor*.

### 2.4 Lignin, cellulose and hemicellulose

#### 2.4.1 Low lignin content

Cellulose, hemicellulose, and lignin are important biopolymers assisting land plants to withstand gravity, grow upward, and conduct water^3, 45^. Lignin in land plants appeared around 400 MYA and was considered an essential evolutionary biomarker during terrestrial adaptation^3^. Lignin content in aquatic algae was first extremely low (3.3%, w/w) and then increased in early land plants such as liverworts (*M. polymorpha*) and mosses (*P. patens*) (4.6%-6.6%, w/w), eventually reaching over 20% in terrestrial angiosperms. However, lignin content in duckweeds is below 7%, much lower than that in other typical angiosperms (Fig. 4a).

**Fig. 4.**
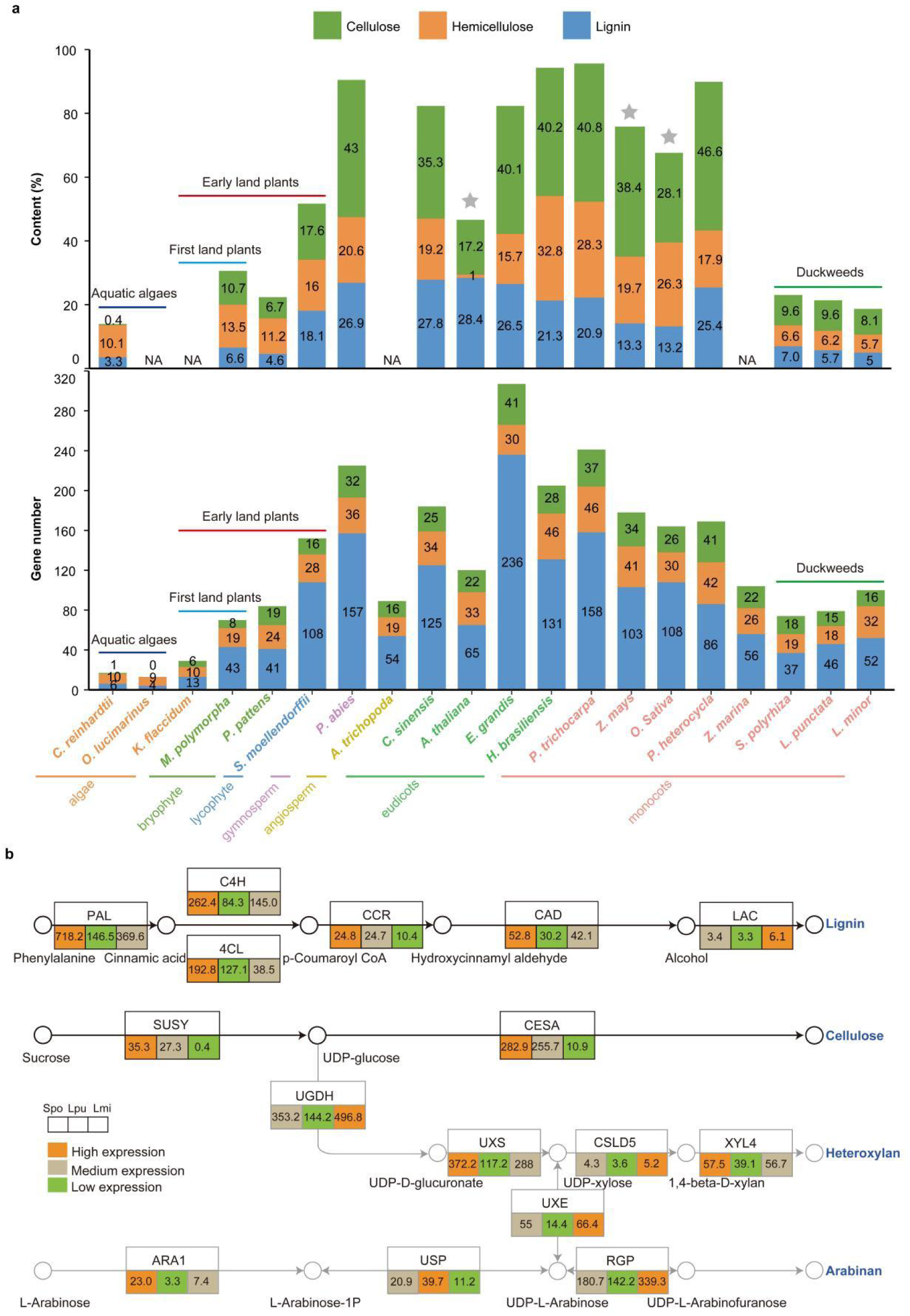
Contents, gene numbers and transcriptional expression of cellulose, hemicellulose, and lignin in duckweeds and other plants. **a**, Contents of cellulose, hemicellulose, lignin, and the number of genes involved in their biosynthesis pathway in duckweeds and other plants (Supplementary Data 4, 5, 6, 18). The numbers in the columns indicate the contents or gene numbers. ★, data available from the stem. NA, no measurement data. **b,** Expression levels of key genes involved in biosynthesis pathway of cellulose, heteroxylan, arabinan, and lignin in duckweeds. The numbers presented in the box are FPKM values of *S. polyrhiza* (Spo), *L. punctata* (Lpu), and *L. minor* (Lmi). Detailed gene information and expression data are provided in Supplementary Data 4, 5, 6.

The number of genes involved in lignin biosynthesis expanded from lower aquatic algae (4-13 genes, eg. *Chlamydomonas* and *Ostreococcus*) to early land plants (e.g. *Marchantia,* 54, and *Physcomitrella,*45) (Fig. 4a). However, these plants lack *F5H*, the key gene for biosynthesis of S lignin^46^, and have only few *CCR*, *CAD,* and *LAC* copies (Supplementary Data 6). This leads to incomplete lignin biosynthesis pathways and therefore low lignin contents (Fig. 4a). On the other hand, duckweed, an embryophyte, has complete lignin biosynthesis pathways that includes *F5H*. Nevertheless, the numbers of *F5H* copies in duckweed is less than that in typical land plants. Despite the difference in their genetic construct, the total number of genes involved in the lignin biosynthetic pathway in duckweed has contracted to a level comparable to that of early land plants (Fig. 4a).

Laccase (LAC) is another important enzyme involved in the final step of lignin synthesis for polymerization^47, 48^. The numbers of *LAC* decreased from 17-74 in typical land plants to 5-8 in duckweed (Supplementary Data 6). In *LAC* family, *LAC4*/*11*/*17* is essential for the formation of lignin, and mutations of those genes lead to different degrees of low lignin content in *Arabidopsis*^48^. All three duckweed species, representing the whole subfamily, lack *LAC4*. Orthologs of both *LAC11* and *LAC17* are detected in *S. polyrhiza*, whereas that of only *LAC11* is found in *L. punctata* and *L. minor* (Extended Data Fig. 5). Nonetheless, expression levels of *LAC11* and *LAC17* in duckweeds is extremely low (Fig. 4b, Supplementary Data 6). The decrease of lignin-related genes and their expression levels among the three species is found consistent with the decreasing trend of lignin content in the order of *S. polyrhiza* (7.0%), *L. punctata* (5.7%), and *L. minor* (5.0%) (Fig. 4a).

Therefore, genomic and transcriptome evidence strongly support the low lignin content of duckweeds and the decreasing lignin content along the evolutionary sequences.

#### 2.4.2 Low cellulose and hemicellulose content

Holocellulose content in duckweeds (13.7-17.0%) was also much lower than that of typical land plants (18.2-73.0%). The number of genes involved in holocellulose biosynthesis in duckweeds (33-48 genes) was contracted compared to that in typical land plants (35-83 genes) (Fig. 4a, Supplementary Data 4, 5). Furthermore, the expression levels of main holocellulose biosynthesis and its precursor genes *SUSY* and *CESA* were consistent with the trends of cellulose content in the three duckweed species (Fig. 4b).

In general, although complete lignocellulose biosynthesis pathway exists in duckweed, the corresponding gene copies contract to levels comparable to that of early land plants (eg. liverworts and mosses). This might be the result of the degeneration of lignocellulose and its function in duckweed to adapt to the new aquatic environment.

### 2.5 Drought stress response

#### 2.5.1 Contraction of LEA genes

Drought is one of the major abiotic stresses that affect the growth of land plants. As a result, various regulation systems against drought stresses have been developed^49, 50^. Late embryogenesis abundant (LEA) proteins are hydrophilic proteins involved in abiotic stress tolerance, especially in drought response^50^. The numbers of *LEA* gene were contracted in duckweeds (13-19 genes) as compared with typical land plants (18-52 genes), especially dehydrins (1-2 genes in duckweed) belonging to LEA protein group II, which is mainly involved in drought resistance (Supplementary Data 7). Moreover, duckweeds lack the orthologous gene of *PvLEA18*, which has also not been found in marine angiosperm *Z. marina* which belongs to the same Order Alismatales as duckweed^9^ (Extended Data Fig.6). This finding implies that aquatic plants no longer need a strong drought resistance ability, resulting in a significant contraction of their drought resistance genes.

### 2.5.2 Contraction of TFs families *bHLH*, *C2H2* and *WRKY*

TFs families expanded drastically during plant terrestrialization. Aquatic algae *Chlamydomonas reinhardtii* possess 208 TFs, first-landing liverwort *M. polymorpha* has 398, and the land dicot *Arabidopsis thaliana* has 1780^40^. In comparison, the three duckweed species have 1076-1148 TFs, positioned between the first-landing liverwort and *A. thaliana*. TFs families of *bHLH*, *C2H2* and *WRKY* play important roles in response to abiotic stresses. These three families contracted significantly in duckweeds, presumably because of the relatively stable aquatic environment (Extended Data Fig.7a). Especially, TFs responding to drought stress in these three families were contracted (*AtWRKY28/57*, *OsWRKY7* and *OsWRKY30*) and even lost (*AtZAT12* and *OsZFP182*) (Extended Data Fig.7b).

The contraction of *LEA* genes and TFs families *bHLH*, *C2H2* and *WRKY* suggested that during the transitional process from terrestrial to aquatic habitats, the regulatory networks in duckweeds related to drought response have declined.

### 2.6 Expanded gene families in duckweed

#### 2.6.1 Expansion of gene families in the flavonoid biosynthesis pathway

Flavonoids play important role in defending plants against bacteria, fungus, and virus^51^. Plant flavonoids originate from charophytic algae, hypothetically closely related to land plants, to adapt to terrestrial environments with more adversity^5, 52^.

K. *flaccidum* has no complete flavonoid biosynthesis pathways, while the gene numbers of these pathways in land model plants *A. thaliana*, *O. sativa,* and *Z. mays* increased significantly and formed complete pathways^40, 53, 54^. Compared with these three model plants, the most prominent expansion pathway in duckweeds is flavonoid biosynthesis, seen markedly expanded in all three species. Flavone & flavonol biosynthesis pathway expands in *S. polyrhiza* and *L. punctata*, and anthocyanin biosynthesis pathway expands in *S. polyrhiza* and *L. minor* (Extended Data Fig. 8). The gene numbers of expanded gene families in flavonoid-related pathways (flavonoid, anthocyanin, and flavone & flavonol biosynthesis) of *S. polyrhiza* (43), *L. punctata* (63), and *L. minor* (33) are much higher than those in land model plants (Supplementary Data 12). Moreover, the flavonoid content in duckweeds was extremely high, reaching 3.51-5.41% (dry basis, d. b.) (Extended Data Fig.9a)^55^, and the anthocyanin content in *Spirodela* genus and *Landoltia* genus were also high (0.22-0.93%, d. b.) (Extended Data Fig. 9b). Both flavonoid and anthocyanin were significantly higher in duckweeds than those in other plants^56–58^. In addition, flavonoids in duckweed exist in a large diversity (more than 30 compounds in *L. punctata*) (Extended Data Fig. 9c).

Flavonoids also assist in maintaining human health, especially in the prevention and treatment of inflammation, heart disease, and cancer^59^. Duckweeds have high flavonoid content and abundant diversity, as well as the expansion of related biosynthetic gene families. Therefore, duckweed has a great potential as a new source of flavonoids^60^. In fact, some human trials have already been carried out with valuable results ^61^.

Notably, the flavonoid and lignin biosynthesis pathways share the same precursor phenylalanine pathway. Compared with land plants, the number of genes in duckweed in the phenylalanine pathway is slightly reduced (Supplementary Data 17). When comparing their downstream pathways, the lignin pathway in duckweed is extremely reduced (Supplementary Data 6), while the flavonoid pathway is significantly expanded (Supplementary Data 12). These sharp changes are possibly adaptations to their new aquatic environment. Meanwhile, flavonoids act as UV “sunscreens” in land plants and is boosted by UV-B induced upregulation^40^. Floating in water with direct exposure from the sun, duckweed doesn’t need much lignin for structural support but requires more flavonoid for stronger antibacterial ability and ultraviolet resistance. This also explains why duckweed can grow vigorously in eutrophic wastewater and can be completely exposed to large amounts of ultraviolet radiation.

#### 2.6.2 Expansion of regulatory genes

As key editing factors, pentatricopeptide repeat (PPR) protein binds to organellar transcripts and influences their expression by altering RNA sequence, turnover, processing, or translation^5, 62^. The two duckweed species, *S. polyrhiza* and *L. punctata,* tend to have much higher numbers of *PPR* (502-579) than those in the three model land plants (407-458) (Supplementary Data 13). This suggests a stronger transcriptional regulation ability in duckweed and explains its flexible environmental adaptability with such small genome size. Transposable element (TE) is associated with the enhancement of regulation ability, adaptability, and diversification^52, 57^. Among the three duckweeds, the proportion of TEs gradually increases along evolutionary direction (13.0% in *S. polyrhiza*, 52.5% in *L. punctata*, and 61.5% in *L. minor*) (Supplementary Table 14). The gradient TEs levels may explain why *Lemna* is more widely distributed than *Spirodela* and *Landoltia* on a global scale.

## 3 Conclusions

Due to the lack of continuous fossil evidence and well-constructed phylogenetic trees that merely relies on a few genes, only limited information on the differentiation of existing species and their ancestors can be excavated in the study of evolution. It is almost impossible to reconstruct the evolutionary process in detail. Herein we studied the extant three species of duckweed and their ancestral fossils. Results demonstrated a progressive evolution through the gradual reduction of several important traits that are most closely related to terrestrial adaptation. This includes stomatal formation and movement, lignin content, root development, phytohormone pathways, and the gradual contraction of their corresponding genes. This study reconstructed the process of how a land plant returns to water, a opposite direction of plant terrestrialization different from textbook definition (Fig. 1a). This finding could deepens and widens our understanding of plant adaptation while shapes our perspective on evolution theories.

## Materials and methods

### Sample preparation and genetic analysis

#### Plant materials

The plants, *Landoltia punctata* strain 0202, *Spirodela polyrhiza* strain 7498, and *Lemna minor* strain 7753 were collected and stored in Chengdu Institute of Biology, Chinese Academy of Sciences (Chengdu, China). They were obtained from Xinjin, China, North Carolina, USA, and Hara, Ethiopia, respectively.

One plant of each stored strain was inoculated to sterilized Hoagland media of 100 ml containing sucrose of 15 g/l at pH 5.0 and cultivated at 25 ± 2 °C with a 16/8 h light/dark cycle and a light intensity of 110 μmol m^-2^ s^-1^ in a climatic chamber. Once duckweed covered the medium surface completely, the plants were transferred to 1/5 Hoagland medium at pH 5.5 and further cultivated for 5 days at 25 ± 2 °C with a 16/8 h light/dark cycle and a light intensity of 110 μmol m^-2^ s^-1^. The cultivated plants were used for the subsequent genomic and transcriptomic sequencing, and biophysiological and biochemical analysis.

### Genome sequencing, assembly, and annotation

Chromosome observation: the number of chromosomes for *Landoltia punctata* was determined by DAPI staining as described in^63^.

Genome sequencing: the genomic DNA of *Landoltia punctata* strain 0202 was extracted using the modified cetyltrimethylammonium bromide (CTAB) method^64^. The DNA concentration and purity were determined using NanoDrop 2000c (Thermo Scientific, Waltham, USA). Short insert paired-end libraries (200 bp, 500 bp, and 800 bp) and larger insert mate-pair libraries (2 Kb, 5 Kb, 10 Kb, and 20 Kb) were constructed and sequenced using Illumina HiSeq 2000 platform at the Beijing Genomics Institute (Shenzhen, China).

Genome assembly: estimation of the genome size of *Landoltia punctata* was based on K-mer frequency distribution analyses. The genome size was calculated according to the following formula:

Genome Size = K_num_/K_depth,_ where K_num_ is the number of K-mer, and K_depth_ is the expected depth of K-mer. The clean reads of 200 bp-, 500 bp-, and 800 bp-insert size were used based on a 17-mer. Prior to assembly, potential sequencing errors of the short insert paired-end libraries were removed or corrected using the k-mer frequency methodology. Genomic assembly was carried out using SOAPdenovo 2.0^65^ with a K-mer of 81 and SSPACE 2.0.2^66^. Gaps in the scaffolds were closed using GapCloser 1.12. Scaffolds less than 150 bp were removed. Scaffolds were screened against NCBI database to eliminate bacterial contaminant. Assembly quality was assessed using BUSCO^67^ and EST methods. We aligned the scaffolds to the genome of *Spirodela polyrhiza* strain 9509^68^ (Sp9509) using LAST (http://last.cbrc.jp/) (e-value ≤ 1.0 × 10^-5^), and the best-scoring match was chosen in cases of multiple matches. Genomic sequences of *Landoltia punctata* were anchored to 20 chromosomes by identification of syntenic blocks between *Landoltia puncatata* and *Spirodela polyrhiza* strain 9509 using ALLMAPS^69^.

#### Genome annotation

Annotation of repetitive sequences: we detected the repetitive sequences of the genome by combining homology-based approach and *de novo* approach. Homology-based approach used TRF^70^, RepeatMasker 4.0.3, and RepeatProteinMask 3 (http://www.repeatmasker.org/) with RepBase^71^. The *de novo* repeats were identified using RepeatModeler 1.0.7 (http://www.repeatmasker.org/RepeatModeler/) and annotated using RepeatMasker 4.0.3^72^. The repetitive sequences predicted by the two approaches were combined and the redundancies were removed.

Gene annotation: *Gene structure prediction*: All repetitive sequences were masked from the genome prior to gene prediction. Gene structure prediction combined homology-based approach, *de novo* approach, and transcript-based approach. For homology-based gene prediction, the target locations of homologous proteins were obtained by aligning the protein sequences of *Arabidopsis thaliana*, *Brachypodium distachyon*, *Zea mays*, *Oryza sativa*, *Sorghum bicolor*, *Lemna minor*, and *Spirodela polyrhiza* to the *Landoltia punctata* strain 0202 genome using TBLASTN^73^ with an e-value of 1 × 10^−5^. For *de novo* gene prediction, AUGUSTUS 2.5.5 (http://augustus.gobics.de/) and Genscan^74^ were applied using gene model parameters trained by *Arabidopsis thaliana*. For transcript-based approach, TopHat 2.1.0^75^ and Cufflinks 2.2.0^76^ were used to assemble transcripts, and the fifth-order Markov model was used to predict ORFs. All available ESTs from *Landoltia punctata* were used to predict genes. Finally, a combined gene set was generated using GLEAN (https://sourceforge.net/projects/glean-gene/) with default parameters, and the redundancies were removed.

*Gene function annotation:* Gene Ontology (GO)^77^ terms and KEGG pathway^78^ identifiers were annotated using an InterProScan analysis^79^. Gene functional descriptions were generated through sequence homology search against InterPro (http://www.ebi.ac.uk/interpro/) and UniProtKB/Swiss-Prot^80^ databases using BLASTP^81^.

### Gene family analyses

Protein sets were collected from 8 species: Klebsormidium flaccidum^41^, Zostera marina^9^, Arabidopsis thaliana^82^, Oryza sativa japonica^83^, Zea mays^84^, Spirodela polyrhiza^85^, Landoltia punctata, and Lemna minor^86^. Following an ‘all-versus-all’ BLASTP (e-value threshold 1 × 10^−3^) comparison, OrthoMCL^87^ (mcl –I 1.5) was used to delineate gene families. Next, gene families involved in root and stomata development, phytohormones pathways, transcription factors, and pentatricopeptide repeat proteins (PPR) were summarized.

For gene family analyses of the lignocellulose biosynthesis pathway, protein sets consisting of 23 species were collected and analyzed using Orthofinder 2.2.1^88^ (Supplementary Table 11).

Predictions and classification of transcription factors of three duckweeds were conducted with iTAK^89^.

### Phylogenetic tree construction and divergence time estimation

For phylogenetic analyses basing on the nuclear genome, the amino acid sequences of 1545 single copy genes shared by the 8 species (*Spirodela polyrhiza*, *Landoltia punctata*, *Lemna minor*, *Oryza sativa*, *Zea mays*, *Arabidopsis thaliana*, *Zostera marina*, and *Klebsormidium flaccidum*) were aligned by Muscle with default settings (Supplementary Data 19). The aligned protein sequences were concatenated into a super gene. Maximum-likelihood (ML) methods for the phylogenetic analyses used PhyML 3.0^90^. Nucleotide substitution model selection was estimated with Smart Model Selection^91^ in PhyML 3.0. JTT**+**G**+**I**+**F was selected as the best-fitting model for ML analyses. One thousand bootstrap replicates were performed to calculate the bootstrap values.

Phylogenetic analyses was based on the chloroplast genome of 23 species (*Chlamydomonas reinhardtii*, *Klebsormidium flaccidum*, *Marchantia paleacea*, *Physcomitrella patens*, *Selaginella moellendorffii*, *Picea abies*, *Amborella trichopoda*, *Glycine max*, *Medicago truncatula*, *Carica papaya*, *Arabidopsis thaliana*, *Vitis vinifera*, *Populus trichocarpa*, *Zea mays*, *Oryza sativa* japonica, *Tofieldia thibetica*, *Symplocarpus*

*renifolius*, *Colocasia esculenta*, *Spirodela polyrhiza*, *Landoltia punctata*, *Lemna minor*, *Wolffiella lingulata*, and *Wolffia australiana*). Data were aligned and concatenated by the HomBlocks pipeline^92^ with default settings (--align -method = Gblocks) (Supplementary Data 20). A tree was constructed using iqtree with the ML method (GTR+F+R4 model) with bootstrap replication of 1000 times^93^. One thousand bootstrap replicates were performed to calculate the bootstrap values of the topology.

The r8s version 1.70 package^94^ was used to infer divergence times on the phylogeny with the fixage command, based on penalized likelihood methods. We set the node divergence time referring to TimeTree database (http://www.timetree.org/).

### Transcriptomic sequencing

Cultivated duckweed fronds were sampled and snap-frozen immediately in liquid nitrogen, then stored at -80 °C prior to RNA extraction. Total RNA was extracted using OMEGA^TM^ Plant DNA/RNA Kit (OMEGA, USA) following the manual. RNA concentrations, quality, and integrity number (RIN) were measured by 2100 Bioanalyzer (Agilent, USA). Construction of cDNA library included mRNAs and ncRNAs isolation, fragmentation, cDNA synthesis, end-repair, A-tailing, adapter ligation, degradation of the second strand, and PCR amplification. Qualified libraries were applied to pair-end sequencing (2×150 bp) using Illumina HiSeq 2500 platform at Gene Denovo Co., Ltd. All raw sequences were evaluated using FastQC v0.11.3. Low quality sequences (reads with adapter, 10% ambiguous base ‘N’, or low quality scores) were filtered out. The high-quality clean reads were aligned to rRNA database via Bowtie2^95^ to remove the rRNA reads. Then, TopHat2^75^ was used to align the reads to the reference genome. Gene expression levels were exhibited as FPKM (fragments per kilobase of transcript per million fragments mapped) values, which were calculated using Cufflinks^76^.

### Physiological and biochemical analysis

#### Morphology observation of duckweeds

Duckweeds cultivated for 3 days were sampled for morphology observation. The morphology of roots and stomata was observed, and the stomata aperture was measured using a Motic BA210 microscope fitted with a Motic imaging accessory (Motic Images Advanced v3.2). Stomata density (SD) and stomata index (SI) were calculated using Eqs. (1) and (2), respectively^96^.

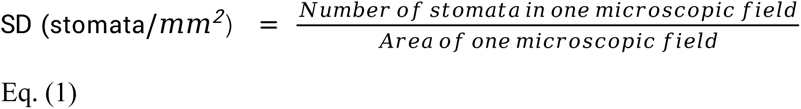

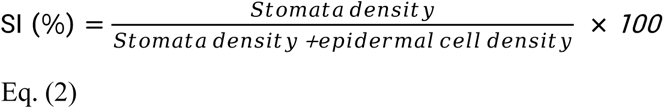

#### Stomatal response of duckweeds to ABA treatment

Stomatal response to ABA treatment referred to the method described by^97^. Briefly, the duckweeds *Spirodela polyrhiza*, *Landoltia punctata*, and *Lemna minor* were cultivated in 1/5 Hoagland media for 3 days. After rinsing with deionized water, the duckweeds were transferred into fresh 1/5 Hoagland medium and placed under white light for 3 hours to induce stomata opening. Then, ABA was added into the medium to a final concentration of 100 μM and treated for 60 min, with treatment in the absence of ABA as control. Stomatal aperture was measured before and after treatment. More than 400 stomata were measured in each group, and at least 10 stomata were measured on each frond.

#### Analytical methods

Lignin and structural carbohydrates, including glucan, xylan, galactan, arabinan, and mannan, were determined according to the method recommended by National Renewable Energy Laboratory, USA^98^. Starch was determined according to the procedure basing on hydrolysis using HCl, as described in our previous study^99^. The difference between glucan and starch was considered cellulose. The sum of xylan, galactan, arabinan, and mannan was considered hemicellulose.

Phytohormones were extracted by a methanol/water (8/2, v/v) solution and determined by an UPLC-ESI-MS/MS system. Fresh duckweed was harvested and snap-frozen in liquid nitrogen, then stored at -80 °C for following analysis. Frozen fresh duckweed of 120 mg was ground and overnight-extracted with 1.2 mL methanol/water (8/2, v/v) at 4 °C. The extract was centrifuged at 12,000 *g* at 4 °C for 15 min. The supernatant was collected and evaporated to dryness under nitrogen gas stream. After re-dissolved in 100 μl 30% (v/v) methanol, the solution was centrifuged, and the supernatant was collected for UPLC-ESI-MS/MS (UPLC, Shim-pack UFLC SHIMADZU CBM30A system, www.shimadzu.com.cn/; MS/MS, Applied Biosystems 6500 Triple Quadrupole, www.appliedbiosystems.com.cn/) analysis.

### Gene family expansion analysis in duckweeds

The gene family expansion analysis in the three duckweeds as compared with model plants was based on the result of OrthoMCL analysis. Expanded families were defined as those where the total number of genes in duckweeds was higher than total number of genes in three model plants. Gene Ontology (GO) terms and KEGG pathways were tested for statistically significantly enrichment using OmicShare tools (http://www.omicshare.com/tools) by conducting a hypergeometric test.

### Identification of SnRK2 (III) in *Lemna minor*

To confirm the precise coding sequence of SnRK2 (III) protein in *Lemna minor* in our study. The SnRK2 (III) coding sequence was cloned for sequencing in *Lemna minor*. Primers were synthesized according to the *Lemna minor* reference genome published by^86^. The SnRK2 (III) annotation number is Lminor_002649 in the reference genome. Forward primer: ATACTGCTCACTCGCCGGGTTC, Reverse primer: ACACGCAGCACGGAGGATTCAA.

The total RNA of *Lemna minor* was extracted using Eastep® Super Total RNA Extraction Kit (Promega, USA) and reverse transcription was performed using GoScriptTM Reverse Transcription System (Promega, USA). Target SnRK2 (III) PCR product was cloned into pClone007 Versatile Simple Vector (TsingKe, China) and used for sequencing. As show in Supplementary Fig. 9, the SnRK2 (III) in *Lemna minor* has the similar coding sequence and length to the SnRK2 (III) in *Spirodela polyrhiza* and *Landoltia punctata*, but the coding sequence of Lminor_002649 gene annotation by^86^ has an additional segment in the intermediate region and lost a segment near 3’ terminus region.

### Geotropism experiment

The duckweeds *Spirodela polyrhiza*, *Landoltia punctata*, and *Lemna minor* were inoculated closely on a filter paper, respectively. The filter paper was infiltrated with 1/5 Hoagland medium. Then, the filter paper was vertically placed along the wall of the beaker into a 500 mL beaker with 450 mL 1/5 Hoagland medium. Keep duckweed 2-3 cm above the liquid level. Duckweed was cultivated for 4 d at 25 °C with a light intensity of 130 μmol photons m^-2^ s^-1^. The morphology of duckweed roots was observed every day.

### Determination of total flavonoids in *Spirodela spp.*, *Lemna spp.* and *Wolffia spp*

The *Spirodela spp.*, *Lemna spp.* and *Wolffia spp.* were cultivated in 50 mL distilled water in 100 mL flask for additional 7 days under a light intensity of 130 μmol photons m^-2^ s^-1^, a photoperiod of 25℃ for 16 h day and 15℃ for 8 h night, 80% relative humidity. And the duckweeds were harvested for biochemical analysis. Three biological replicates were established independently. The total flavonoid content was determined by HPLC (Thermo spectra system AS3000, USA)-UV (Thermo UV6000 Detector, USA) referring to the method on literature^55^.

### Determination of anthocyanins in *Spirodela* and *Landoltia*

The *Spirodela polyrhiza* (7 ecotypes), *Spirodela intermedia* (3 ecotypes) and *Landoltia punctata* (2 ecotypes) were cultivated in 250 mL 1/5 Hoagland’s medium, light intensity of 110 μmol m^-2^s^-1^. Briefly, for determination of anthocyanins, 50 mg dry powder of duckweeds were dissolved in 5 mL 70% methyl alcohol (containing 2% methanoic acid) and ultrasonicated for 10 min, then standing for 5 hours in dark for extraction of anthocyanins. The crude extracts were used for HPLC analysis (Thermo spectra system AS3000 and UV6000 Detector, USA) after filtered by 0.22 μm filter. The elution time was 15 min, mobile phase A was 10% methanoic acid, B was 100% methyl alcohol, flow rate was 0.6 mL/min. determination wavelength was 530 nm, scanned area was 200-798 nm. The cornflower-3-glucoside was used as standard for determination of anthocyanins in duckweeds.

### Determination of variety of flavonoids in *Landoltia punctata*

Plant powder (0.25 g) was ultrasonically extracted with 5.0 mL of 70% MeOH for three times (30 min for each time). The solvent was concentrated to dryness in a rotary evaporator at 60 °C under reduced pressure. The residue was dissolved in 1.0 mL 70% methanol in a volumetric flask and then filtered through 0.2 μm PTFE membrane by a syringe filter before use. Analytical high-performance liquid chromatography (HPLC) of flavonoid profile of the crude extracts were carried out on a LC CO-100 instrument equipped with an UV6000 DAD detector (Thermo Electron Corporation), AXW-8 temperature controller, AS3000 auto sampler and Kromasil 100-5C18 and UniSil®10um-100 C18 columns (250 × 4.6 mm) at 326 nm. The UPLC-ESI-QTOF-MS2 was performed by using Waters Vion® IMSQ Tof system equipped with a photodiode-array detector PDAeλ^100^. Same condition was used for mass spectrometers equipped with a Kromasil 100-5C18 column as above. High purity nitrogen (N_2_) was used as nebulising gas, while ultra-high purity helium (He) was used as the collision gas. The ion source was operated in the negative and positive modes. The mass scan arrange was set as m/z 50-1000 for TOF MS2 scan. The following parameter conditions were used: ion spray voltages, 3000 v in positive and 2500 v in negative mode, source temperature 120 °C, desolvation temperature 450 °C, cone gas 50 L/h, desolvation gas 800 L/h and scan time 0.200 s. The mobile phase and condition were same as described above.

### Statistical analysis

Wilcoxon rank-sum test was applied to test for significant differences of stomatal response to ABA treatment experiments, more than 400 stomata were measured in each group, and at least 10 stomata were measured on each frond.

Gene Ontology (GO) terms and Kyoto Encyclopedia of Genes and Genomes (KEGG) pathways those were statistically significantly enriched in these expansion families were tested using OmicShare tools (http://www.omicshare.com/tools) by conducting a hypergeometric test.

## Acknowledgments

We thank Zhongyan Wang for technical support. We thank Wan Xiong for language editing. We also thank Ping Mao for providing information. This research was supported by Innovation Academy for Seed Design, CAS; National Aquatic Biological Resource Center (NABRC); the National Natural Science for General Foundation of China (31770395); Key deployment projects of Chinese Academy of Sciences (ZDRW-ZS-2017-2-1); and Biological Resources Programme, Chinese Academy of Sciences (KFJ-BRP-008).

## Author contributions

H.Z, Y.F and Y.J conceived and designed the study, H.Z, Y.F and Y.J supervised and funded the study, Y.F, X.T, Y.J, A.D, Z.L, L.G, Y.Xiao, Y.Xu, S.C, L.T, F.Z, G.L, Jinmeng.L, L.X and X.H.C performed the experiments, X.T, A.D, Y.D, Y.Z, Y.C, Z.Y and L.C performed bioinformatics analysis, H.Z, Y.F, X.T, A.D, Y.D, Z.L, K.H, Y.Z, L.G, Y.Xiao, Y.Xu, S.C, L.T, S.W, Jiatang.L, Z.Y, L.C, F.Z, N.W, Y.Z, H.C and Q.Z contributed to the discussion and important intellectual content, Y.F, H.Z, X.T, A.D, Z.L, K.H, S.C, S.W, Jiatang.L, L.Z, Y.Z, and Q.Z wrote and revised the manuscript.

## Data and materials availability

All data are available in the manuscript, the supplementary or at publicly accessible repositories. Raw reads of whole genome sequencing of *Landoltia punctata* 0202 have been deposited at NCBI under BioProject ID: PRJNA546087 and the whole genome data of other species used in this study are available in supplementary materials. All transcriptomes have been uploaded to NCBI under BioProject ID: PRJNA670783, PRJNA670784 and PRJNA670786. The duckweeds are available from the Duckweed Resource Bank in Chengdu Institute of Biology, Chinese Academy of Sciences.

---

**Extended Data Fig.1.**
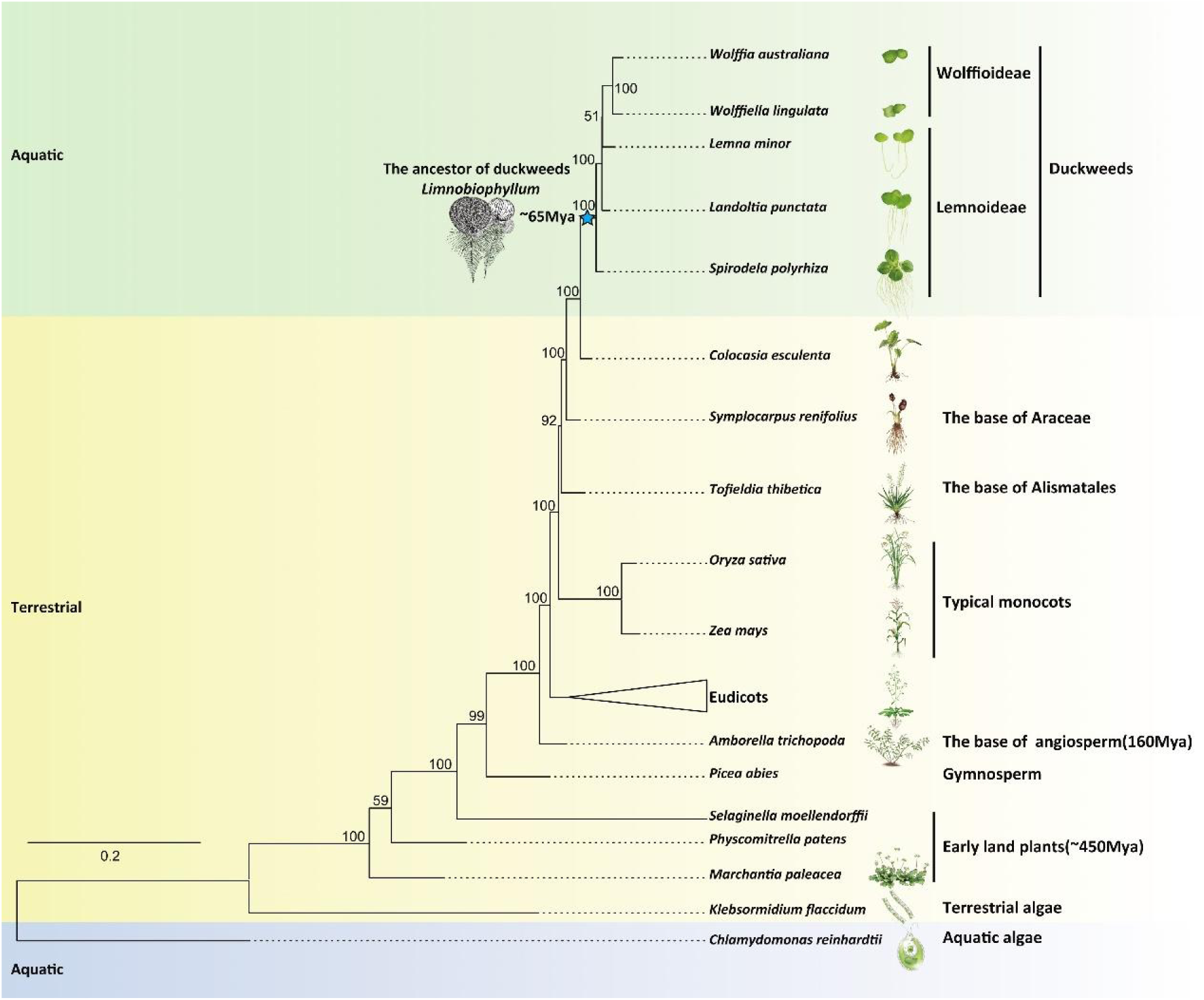
Phylogenetic analysis based on chloroplast genomes from algae (blue), terrestrial plants (yellow), and aquatic plants (green). Numbers at the nodes are bootstrap values. Tree was rooted using *Chlamydomonas reinhardtii* as the outgroup taxon. The eudicots included *Glycine max*, *Medicago truncatula*, *Carica papaya*, *Arabidopsis thaliana*, *Vitis vinifera*, and *Populus trichocarpa*. ★, the ancestor of duckweeds (*Limnobiophyllum scutatum*).

**Extended Data Fig.2.**
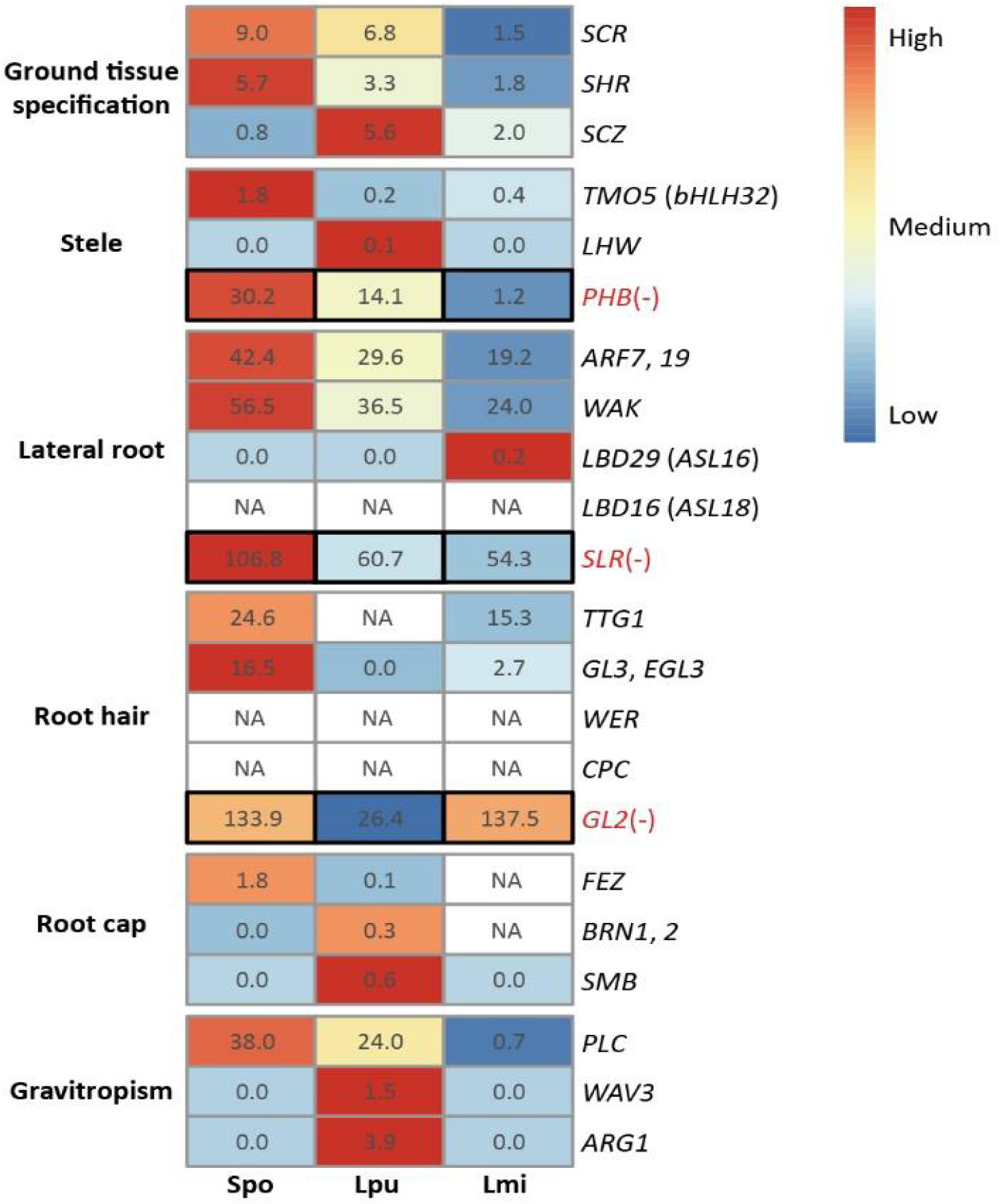
Expression levels of genes involved in root development in duckweeds. Numbers in the boxes are FPKM values. Spo, *Spirodela polyrhiza*; Lpu, *Landoltia punctata*; Lmi, *Lemna minor*; black boxes, negatively related to root development; NA, genes lost. Detailed gene names are provided in Supplementary Data 1.

**Extended Data Fig.3.**
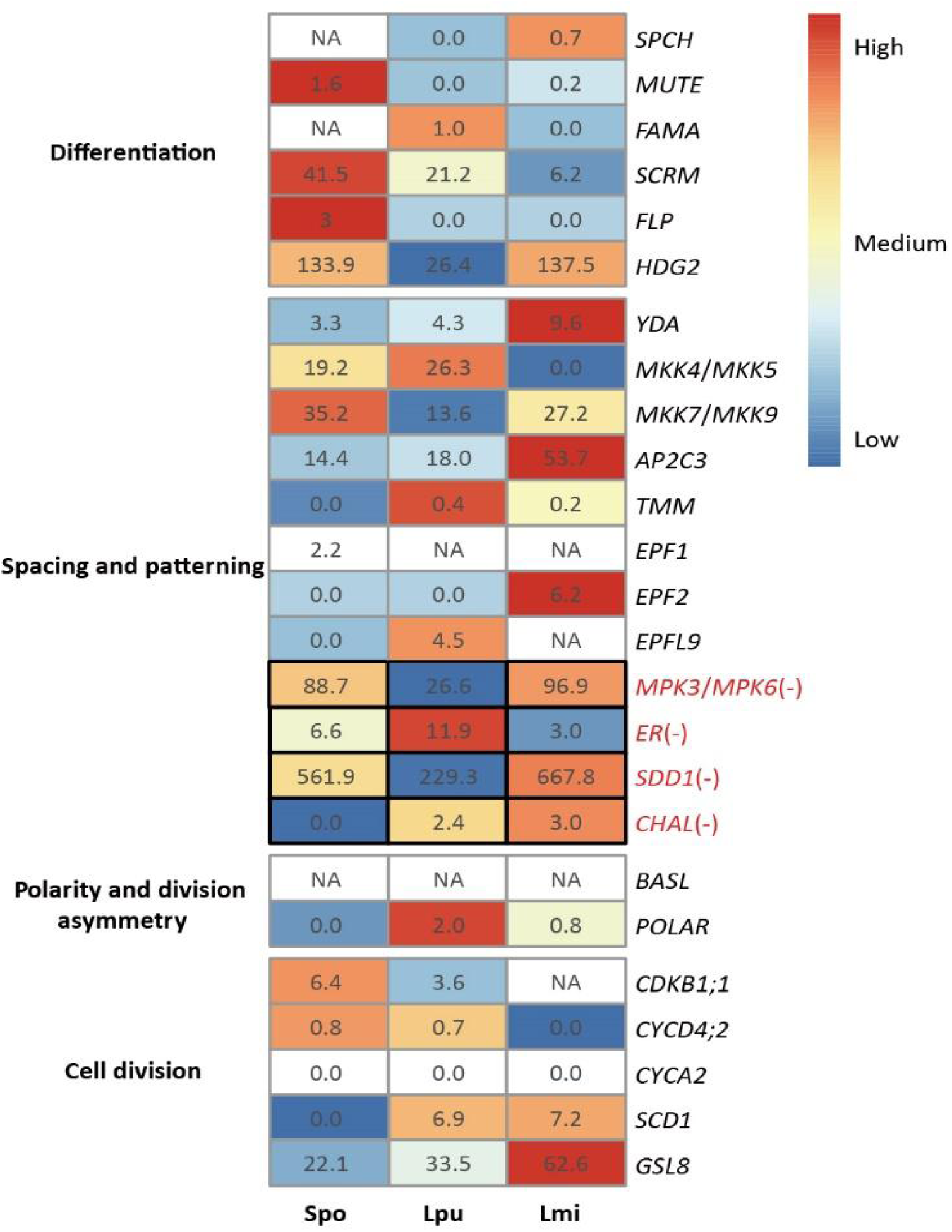
Expression levels of genes involved in stomata development in duckweeds. Numbers in the boxes are FPKM values. Spo, *Spirodela polyrhiza*; Lpu, *Landoltia punctata*; Lmi, *Lemna minor*; black boxes, not related to stomata development; NA, genes lost. Detailed gene names are provided in Supplementary Data 2.

**Extended Data Fig.4.**
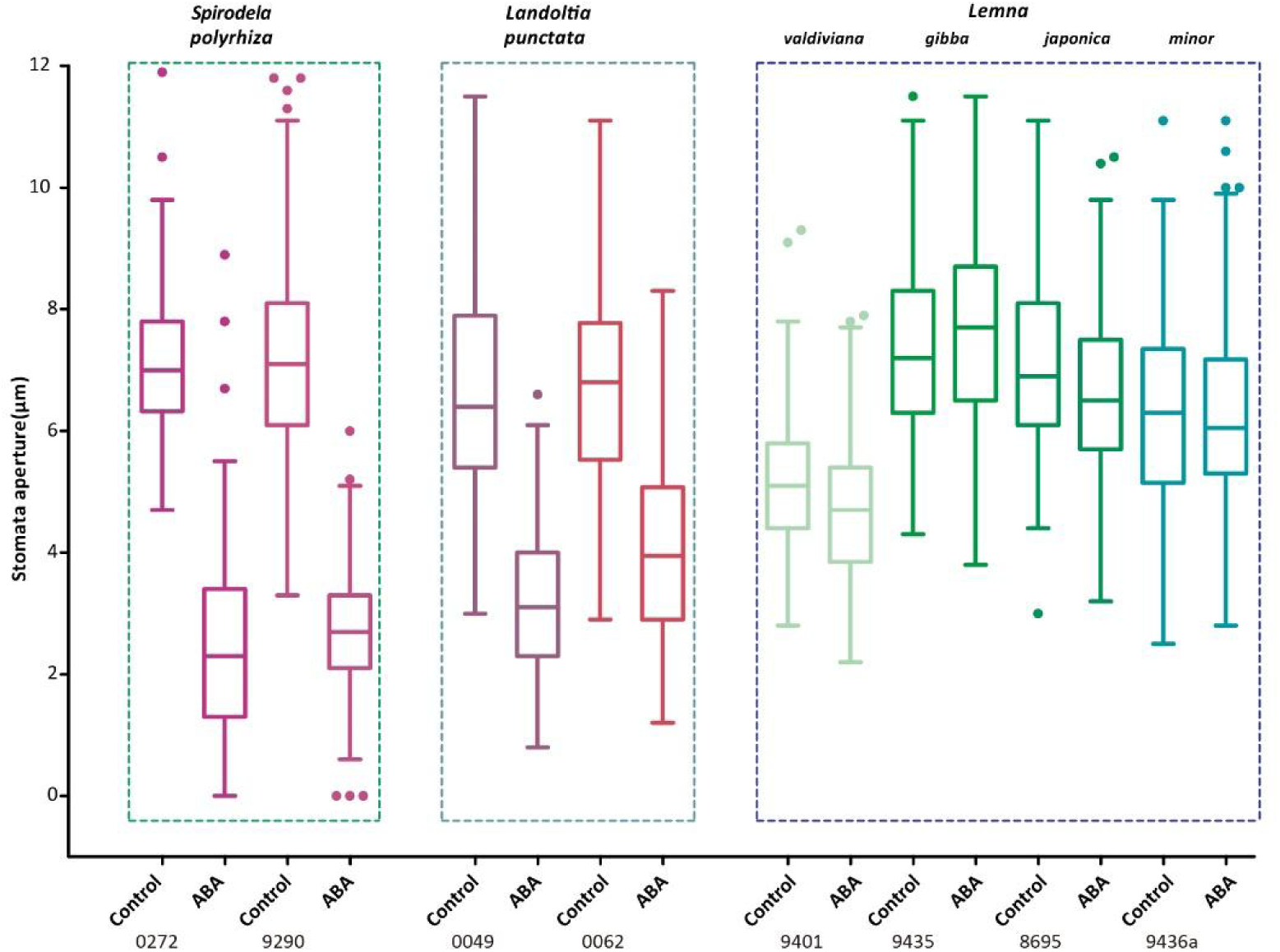
Distribution of stomatal aperture in duckweeds under ABA treatment. Duckweeds were placed in 1/5 Hoagland for 3 h under white light to induce stomata opening and further incubated in the absence (control groups) or in the presence 100 μM ABA (ABA groups).

**Extended Data Fig.5.**
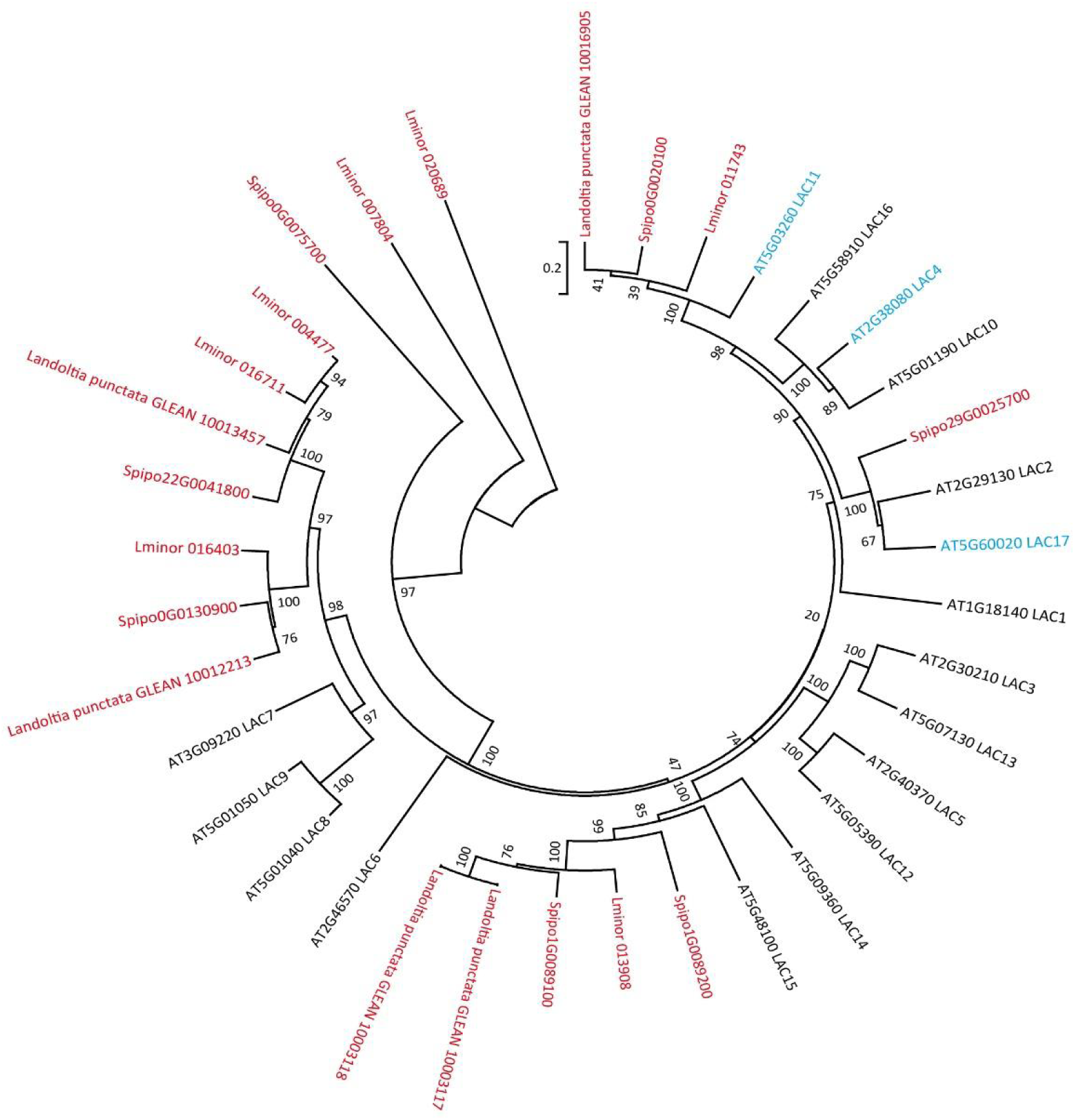
Phylogenetic analyses of laccase (LAC) genes in *Arabidopsis thaliana* and duckweeds. Key genes involved in lignin synthesis of *Arabidopsis thaliana* (*AtLAC4*, *AtLAC11* and *AtLAC17*)^47, 48^ highlighted in red. One homologous gene of *AtLAC11* occurs in each duckweed genome, one *AtLAC11* homologue in *Spirodela polyrhiza*, and no *AtLAC4* homologues in duckweeds.

**Extended Data Fig.6.**
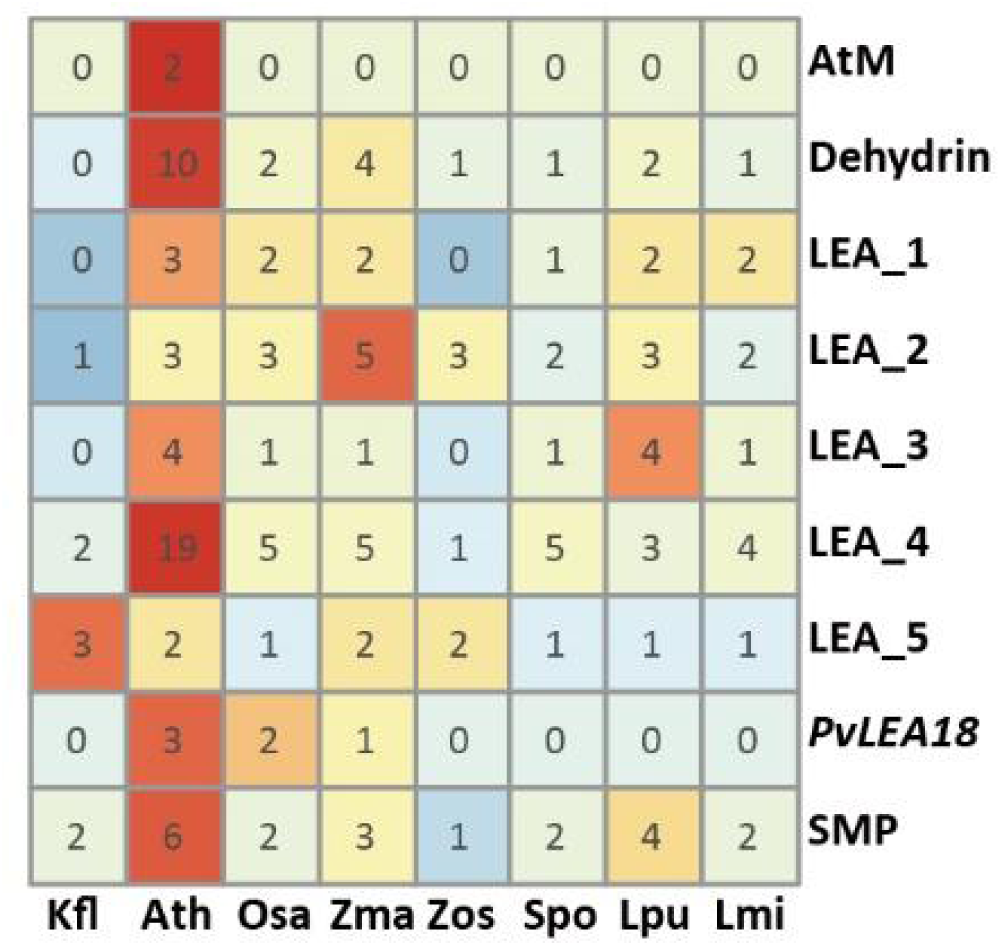
Numbers of late embryogenesis abundant (LEA) protein genes in duckweeds and terrestrial plants. Kfl, *Klebsormidium flaccidum*; Zos, *Zostera marina*; Spo, *Spirodela polyrhiza*; Lpu, *Landoltia punctata*; Lmi, *Lemna minor*; Ath, *Arabidopsis thaliana*; Osa, *Oryza sativa*; Zma, *Zea mays*. Detailed gene information provided in Supplementary Data 7.

**Extended Data Fig.7.**
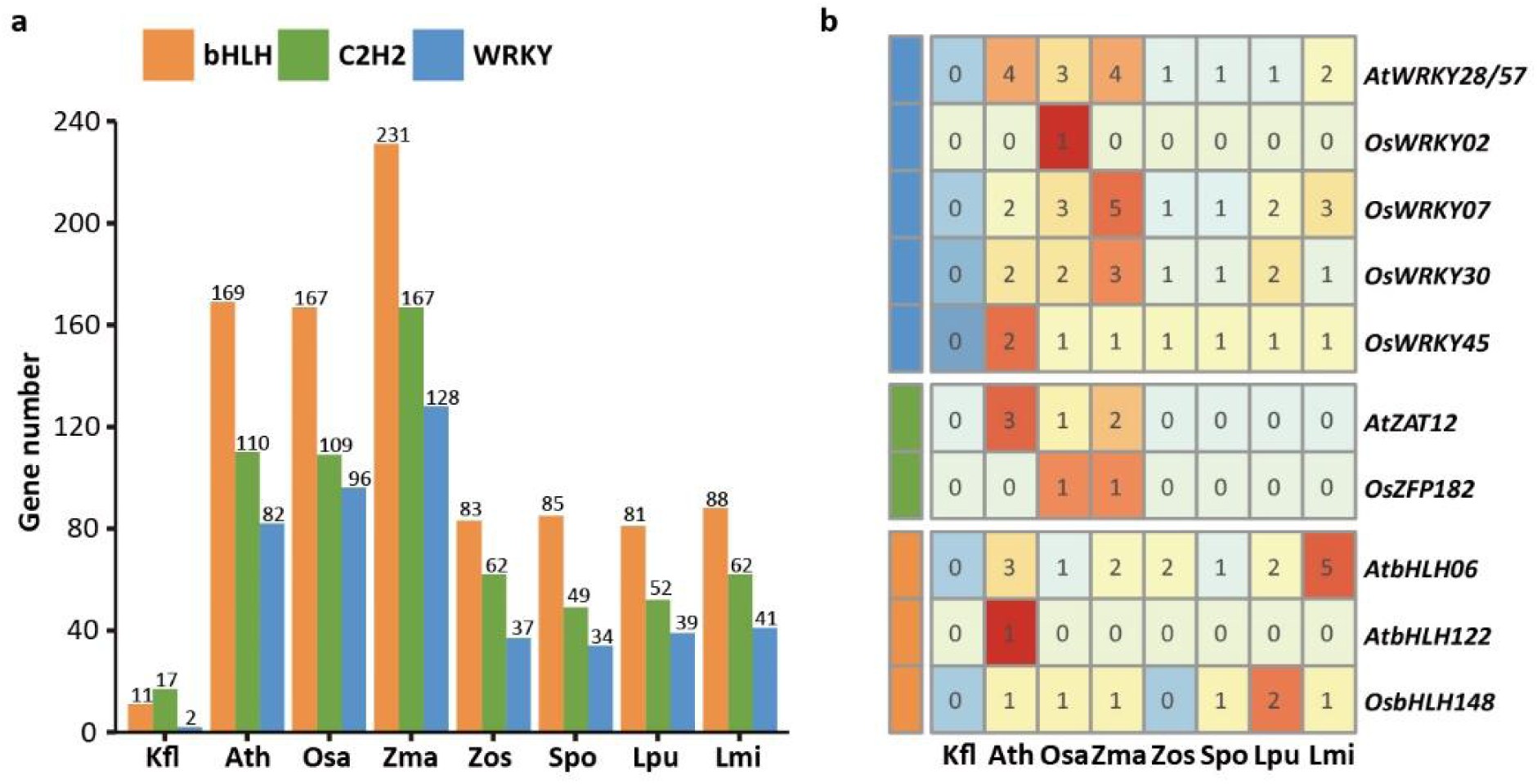
Numbers of transcription factor (TF) genes involved in stress response in duckweeds and land plants. **a**, Number of TF genes in the families bHLH, C2H2, and WRKY. Detailed gene information was provided in Supplementary Data 9c, d, e. **b**, Number of TF genes involved in the drought stress signaling network. Kfl, *Klebsormidium flaccidum*; Zos, *Zostera marina*; Spo, *Spirodela polyrhiza*; Lpu, *Landoltia punctata*; Lmi, *Lemna minor*; Ath, *Arabidopsis thaliana*; Osa, *Oryza sativa*; Zma, *Zea mays*. Detailed gene information was provided in Supplementary Data 9a.

**Extended Data Fig.8.**
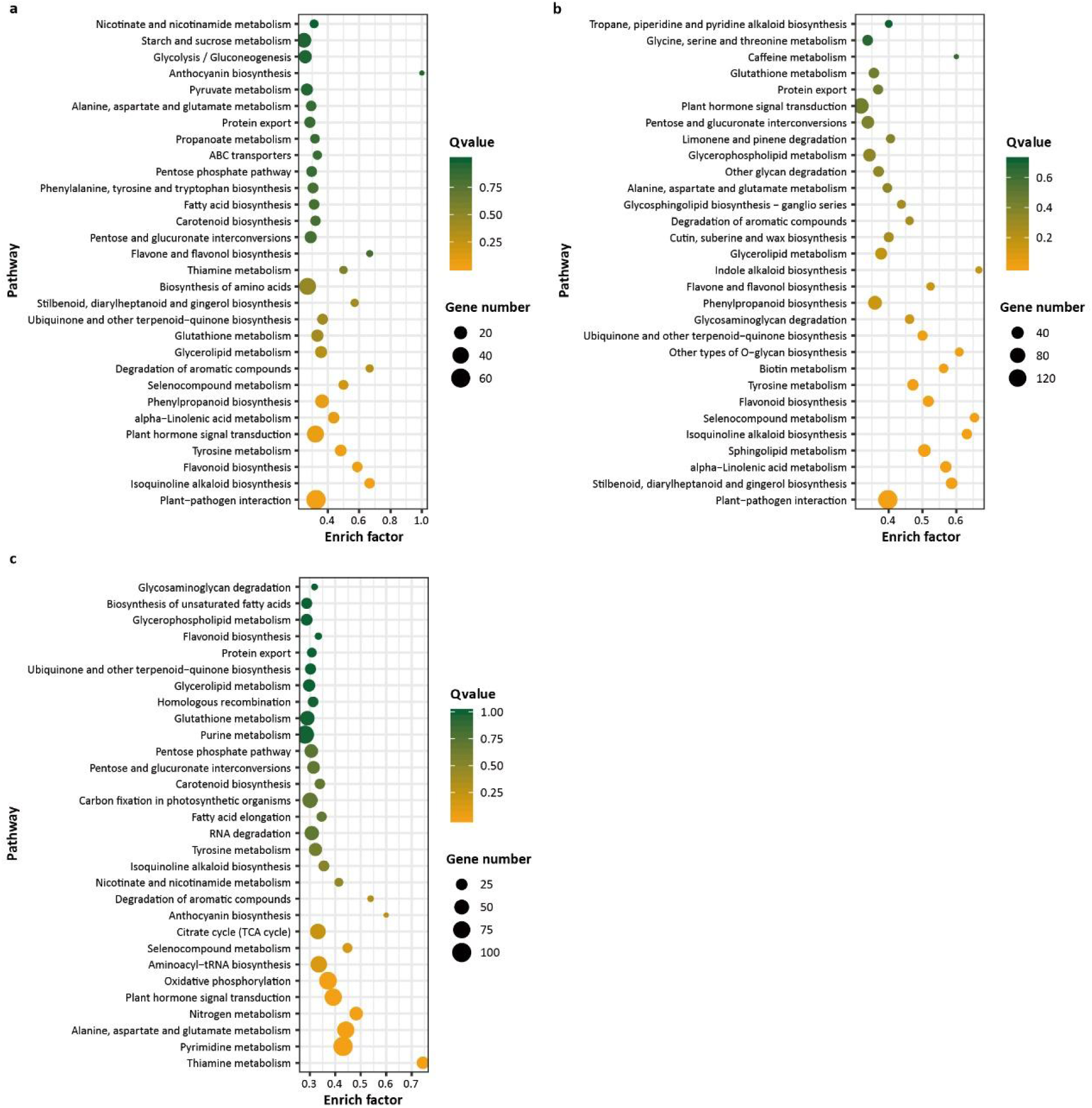
Enriched top 30 KEGG pathways of expanded gene families in duckweeds as compared with model plants *Arabidopsis thaliana*, *Oryza sativa*, and *Zea mays*. **a**, Enriched top 30 pathways in *Spirodela polyrhiza*. **b**, Enriched top 30 pathways in *Landoltia punctata*. **c**, Enriched top 30 pathways in *Lemna minor*. Detail information provided in Supplementary Data 12. The flavonoid, anthocyanin, flavone and flavonol biosynthesis pathways are enriched in duckweeds, which is consistent with the high flavonoid content of duckweeds (Extended Data Fig.9a)^101^.

**Extended Data Fig.9.**
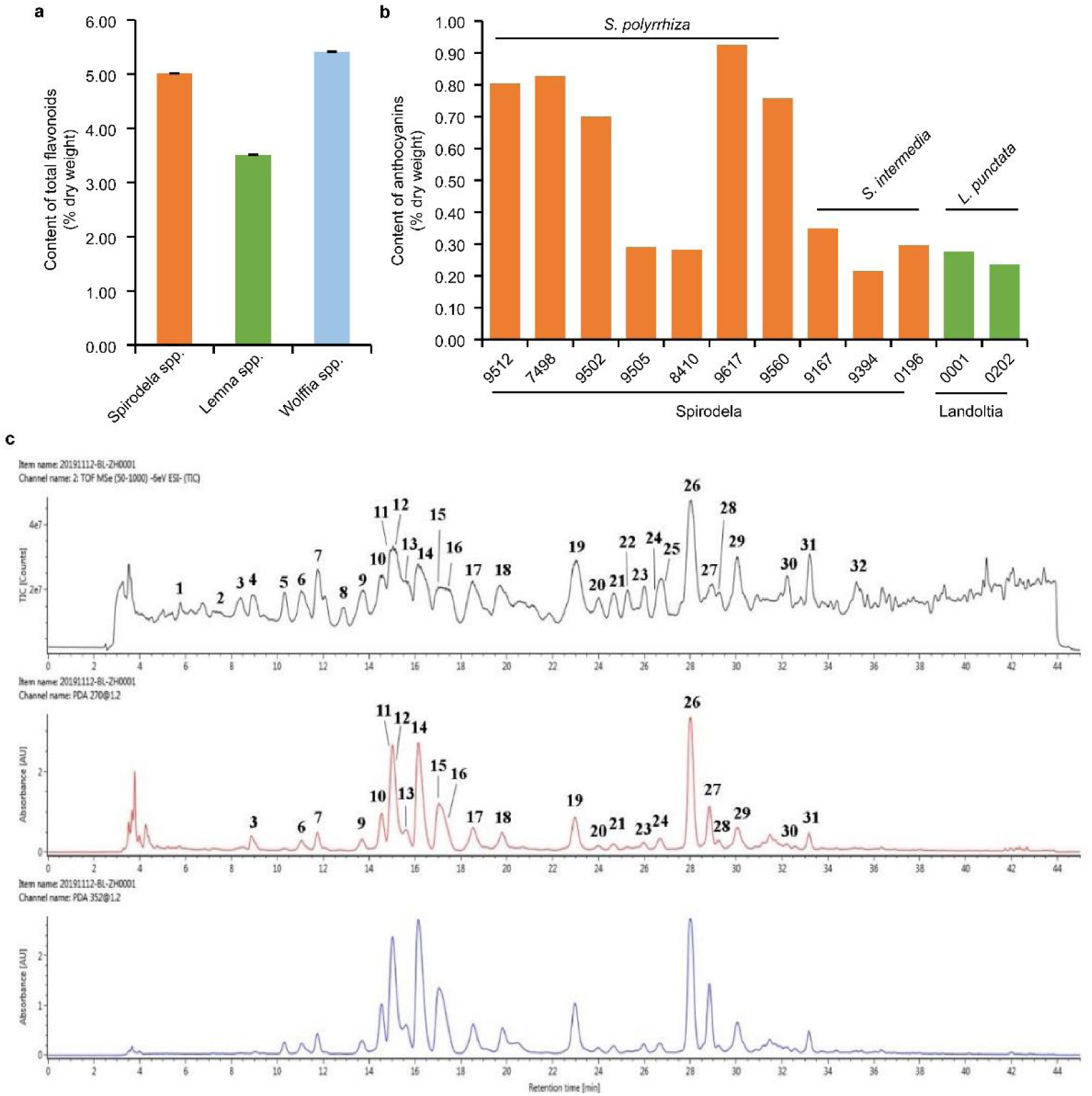
The content and variety of flavonoids in duckweeds. **a,** The content of total flavonoids in *Spirodela spp.*, *Lemna spp.* and *Wolffia spp.*. Three biological replicates were performed independently. The average content was showed and the error bars show the SD value. **b,** The content of anthocyanins in *Spirodela polyrhiza* (7 ecotypes), *Spirodela intermedia* (3 ecotypes) and *Landoltia punctata* (2 ecotypes). **c,** The HPLC-MS and HPLC-TIC chromatograms of *Landoltia punctata* in negative ion mode. Upper panel, middle panel and lower panel show the HPLC-TIC, HPLC-MS and HPLC chromatograms, respectively. There are more than 30 obvious peaks observed in both HPLC-MS and HPLC-TIC chromatograms, indicating more than 30 flavonoids in *Landoltia punctata*.

**Extended Data Table 1.**
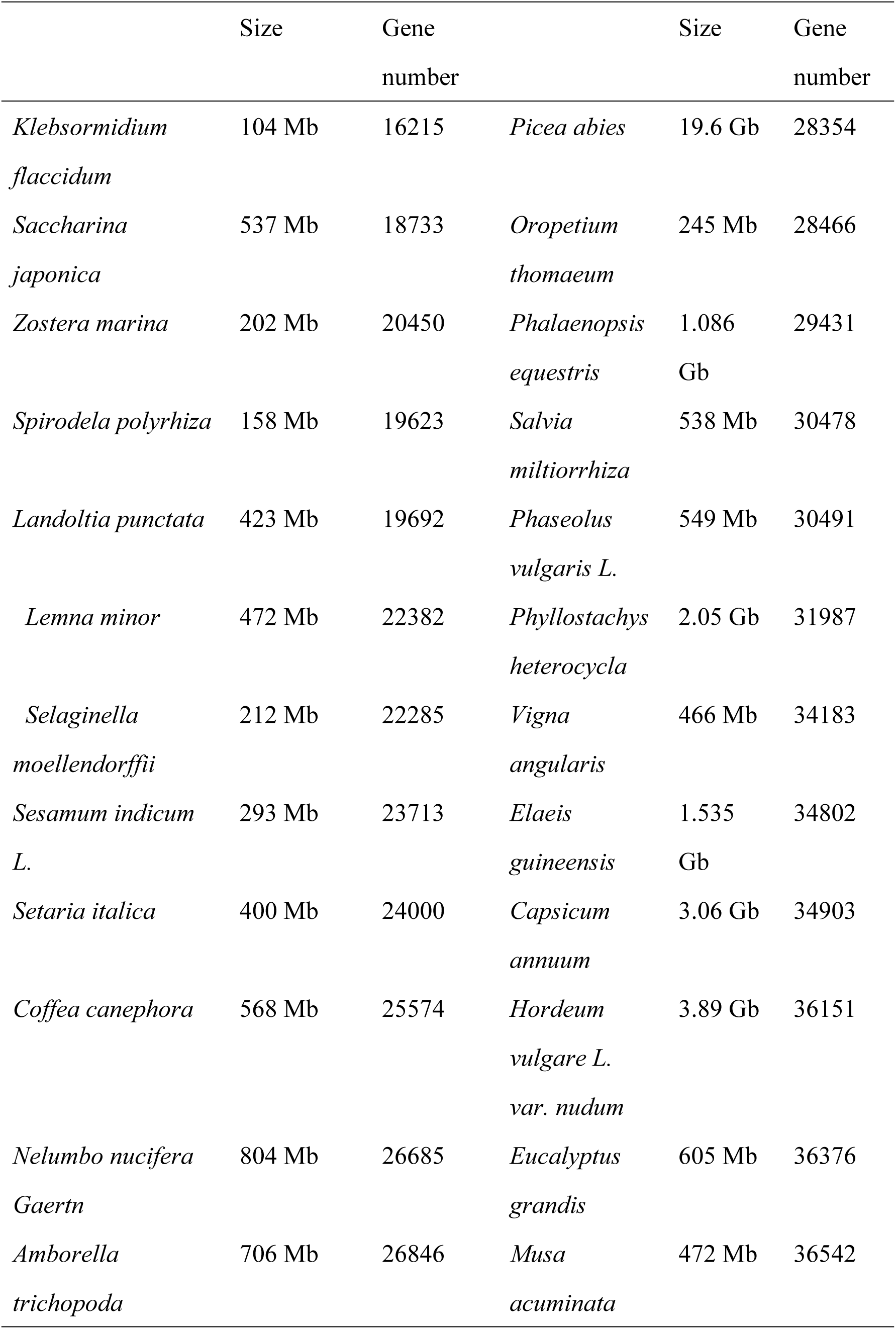
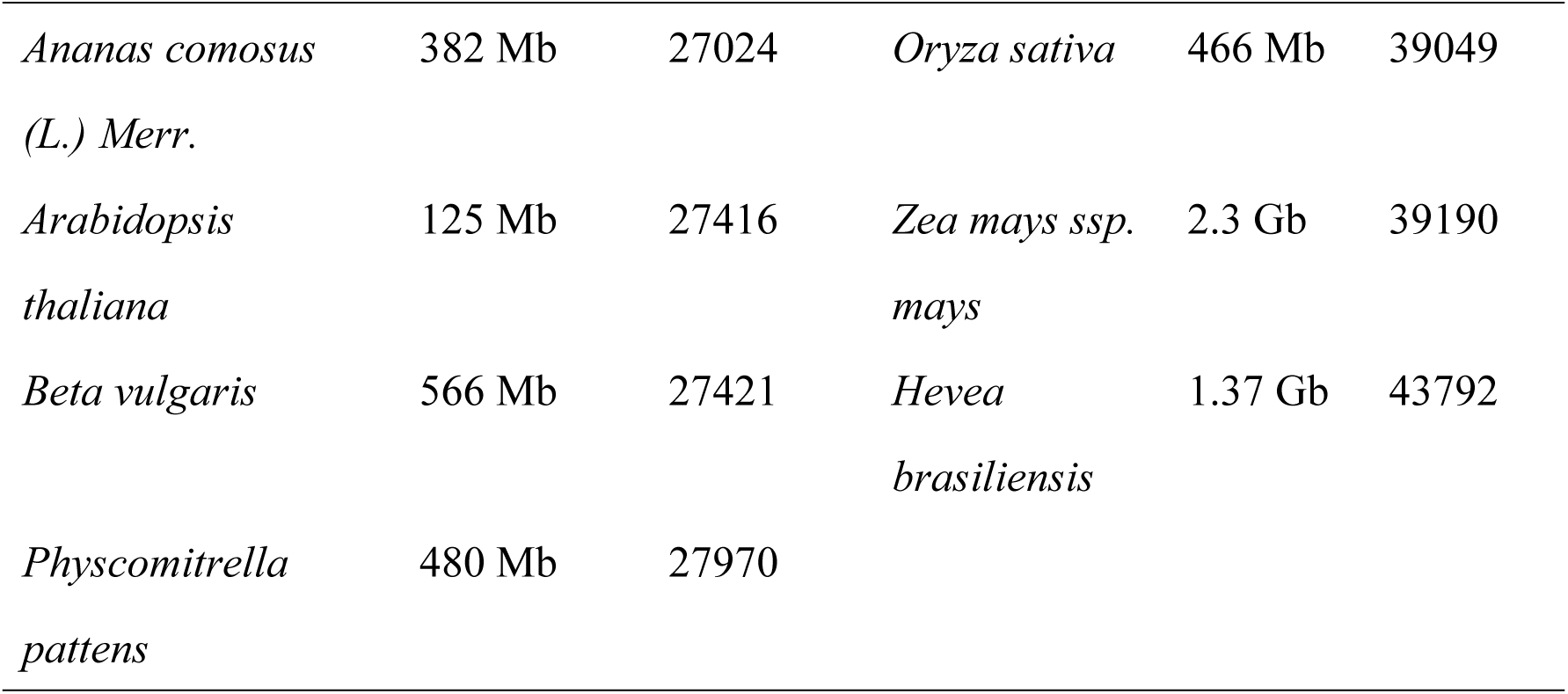
The size and gene number of published plants genomes.

