## Supplementary Materials for "Duckweed evolution: from land back to water"

**This file includes:**

Supplementary Text

Supplementary Figures 1-9

Supplementary Tables 1 to 14

Captions for Supplementary Data 1 to 20

**Other Supplementary Materials for this manuscript include the following:**

Supplementary Data 1 to 18 (.xlsx)

Supplementary Data 19 (.txt)

Supplementary Data 20 (.txt)

### Supplementary Text

1 Chromosomes of *Landoltia punctata*

Metaphase spreads indicated that *Landoltia punctata* strain 0202 has 2n=40 chromosome (Supplementary Fig. 1), the same as *Spirodela polyrhiza* strain 7498 and *Lemna minor* strain 5500.

2 Estimation of the *Landoltia punctata* genome size

Sequencing generated 166.4 Gb of raw reads. After removing adapter sequences, low-quality reads, and duplicate reads, 96.6 Gb of clean reads remained (Supplementary Table 1). More than 80% of bases had sequencing depths of > 100×. Bases with sequencing depth of < 10× accounted for approximately 2.4%. The average sequencing depth was 224.4×

(Supplementary Fig. 2). The sequencing depth was sufficient for subsequent genome assembly. The estimated genome size of *Landoltia punctata* was approximately 415 Mb (Supplementary Fig. 2 and Supplementary Table 2).

3 Chromosome anchoring

Supplementary Fig. 5 shows that gene homology is conserved between the two species. In total, 403.5 Mb (95.5% of *Landoltia punctata* genome) of the sequences of *Landoltia punctata* anchored to chromosomes. The remaining scaffolds containing approximately 18.9 M were combined as chromosome 0.

4 Summary of assembly and annotation statistics of *Landoltia punctata*

The final assembly consisted of 48,966 scaffolds (63,986 contigs), covering 422.4 Mb of the genome with a scaffold L50 of 4.0 Mb and corresponding contig L50 of 54.0 kb; these summaries were longer than most of plant assemblies applying next generation sequencing (Supplementary Table 3 and 4). The GC content of *Landoltia punctata* is 36.5% (Supplementary Table 3).

Analyses predicted 19,692 protein-coding genes in *Landoltia punctata* whose length averaged 3,586 bp, with an average of 5 exons. The functions of 15,890 predicted genes (80.7%) were identified (Supplementary Table 9 and 10). Repetitive sequences accounted for 59.2% of *Landoltia punctata* genome (Supplementary Table 7), which was higher than that of *Spirodela polyrhiza* (*Spirodela polyrhiza* strain 7498, 14.7%; *Spirodela polyrhiza* strain 9509, ~25.3%). In the genome of *L. punctata*, long terminal repeats (LTRs) were the most abundant retrotransposon, accounting for 19.4% of the assembly (Supplementary Table 8).

Quality of assembled genome and annotation was assessed by BUSCO and/or EST. For the genome assembly evaluation, we compared the genome assembly quality of *Landoltia punctata* with those of the model plants and species related to duckweed. BUSCO assessment, including 303 BUSCOs, showed that 88.8% of the set of core eukaryotic genes were present (248 complete single-copy and 21 complete duplicated). The numbers of fragmented and missing BUSCOs were 7 and 27, respectively. Our assembly had more completely represented BUSCOs than all of the model plants, except *Arabidopsis* (Supplementary Fig. 4). EST assessment showed that 3,720 ESTs out of 3,794 (98.0%) were mapped to the assembly (Supplementary Table 5 and 6). This indicated a high quality of genome assembly. For genome annotation evaluation, among 303 BUSCOs, 95.1% of the set of core eukaryotic genes were present completely (261 complete single-copy and 27 complete duplicated). The numbers of fragmented and missing BUSCOs were 11 and 4, respectively. Compared with the assessments of relative species, the most BUSCOs of *L. punctata* were detected completely, indicating high quality of annotation in this study (Supplementary Fig. 6).

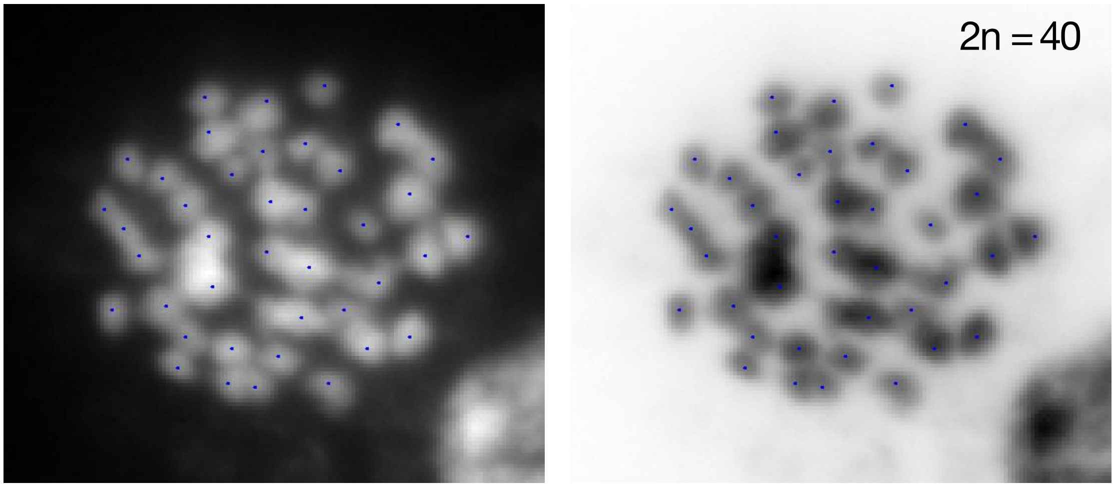


**Supplementary Fig. 1.**

Metaphase chromosomes of *Landoltia punctata* (2n=40).

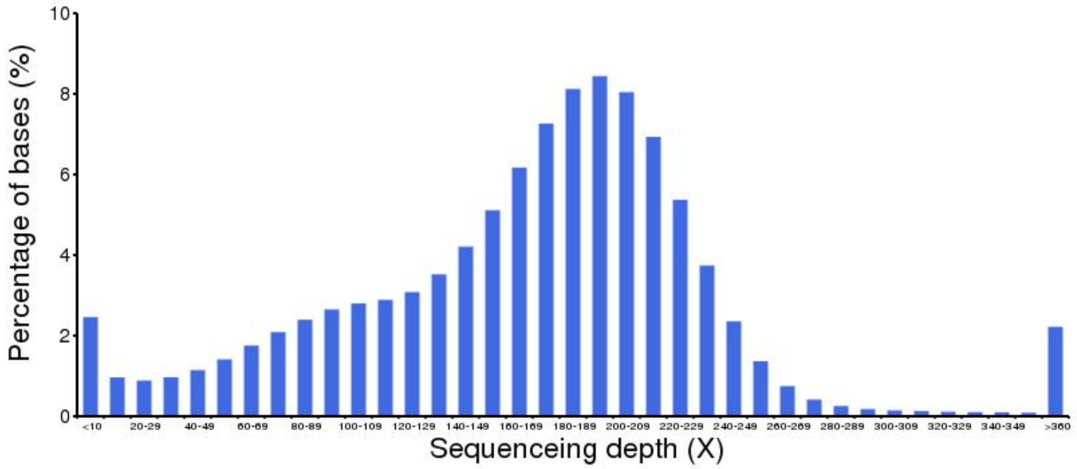


**Supplementary Fig. 2.**

Genomic sequencing depth.

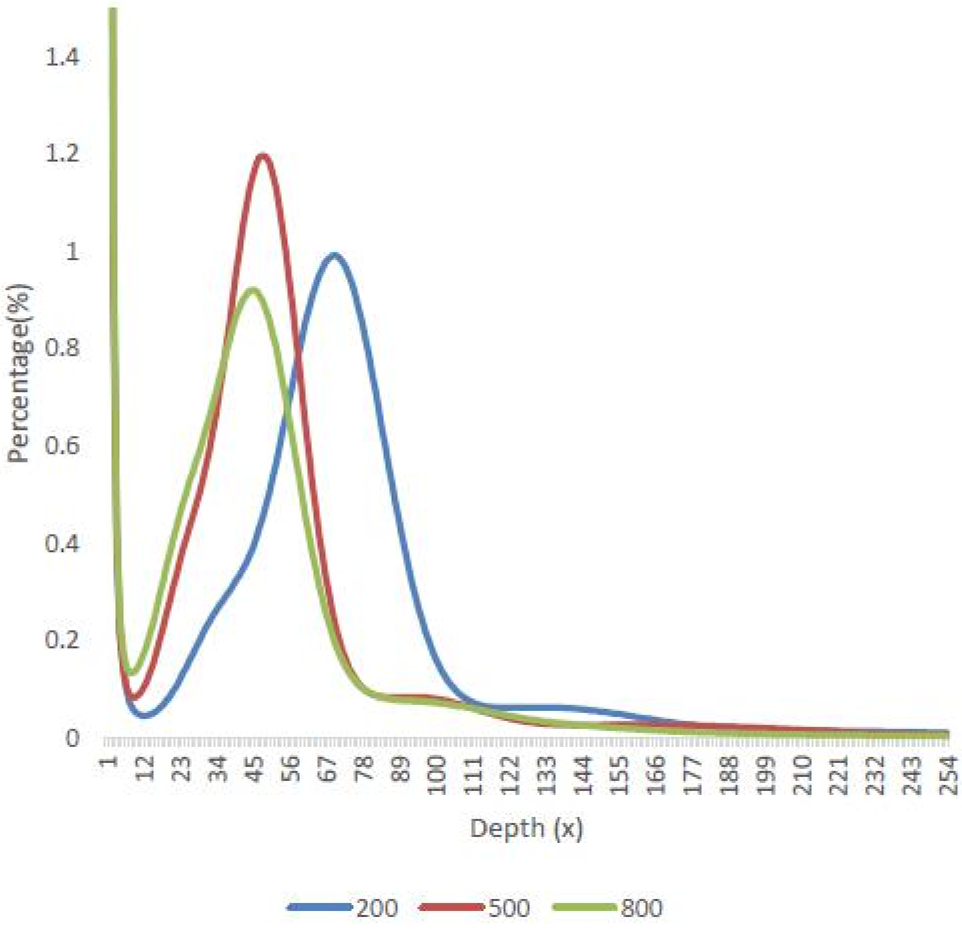


**Supplementary Fig. 3.**

K-mer frequency distribution based on a 17-mer.

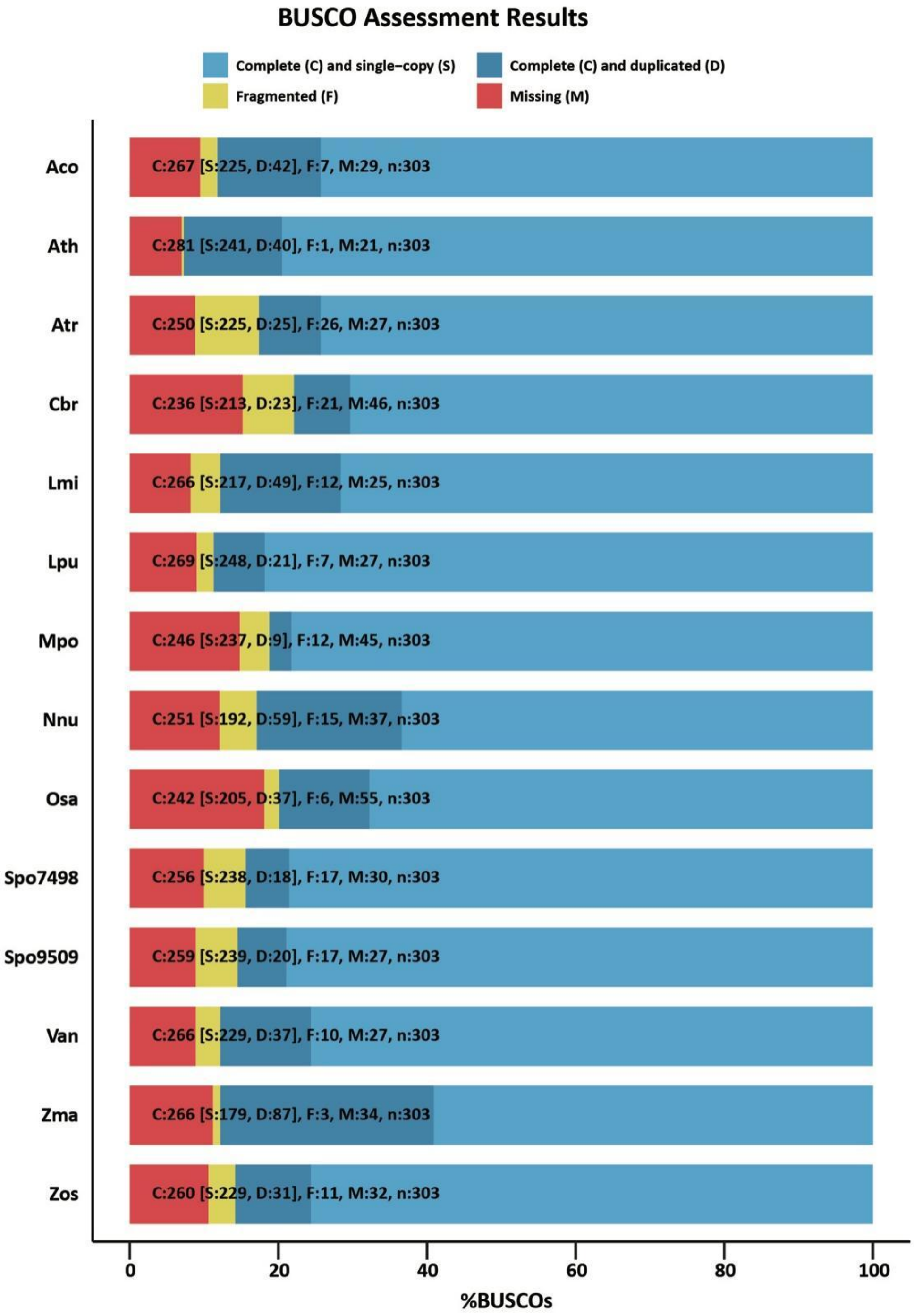


**Supplementary Fig. 4.**

Assessment of genome assembly using BUSCO.

Aco, *Ananas comosus* (L.); Ath, *Arabidopsis thaliana*; Atr, *Amborella trichopoda*; Cbr, *Chara braunii*; Lmi, *Lemna minor*; Lpu, *Landoltia punctata*; Mpo, *Marchantia polymorpha*; Nnu,

*Nelumbo nucifera*; Osa, *Oryza sativa*; Spo7498, *Spirodela polyrhiza* strain 7498; Spo9509, *Spirodela polyrhiza* strain 9509; Van, *Vigna angularis*; Zma, *Zea mays*; Zos, *Zostera marina*.

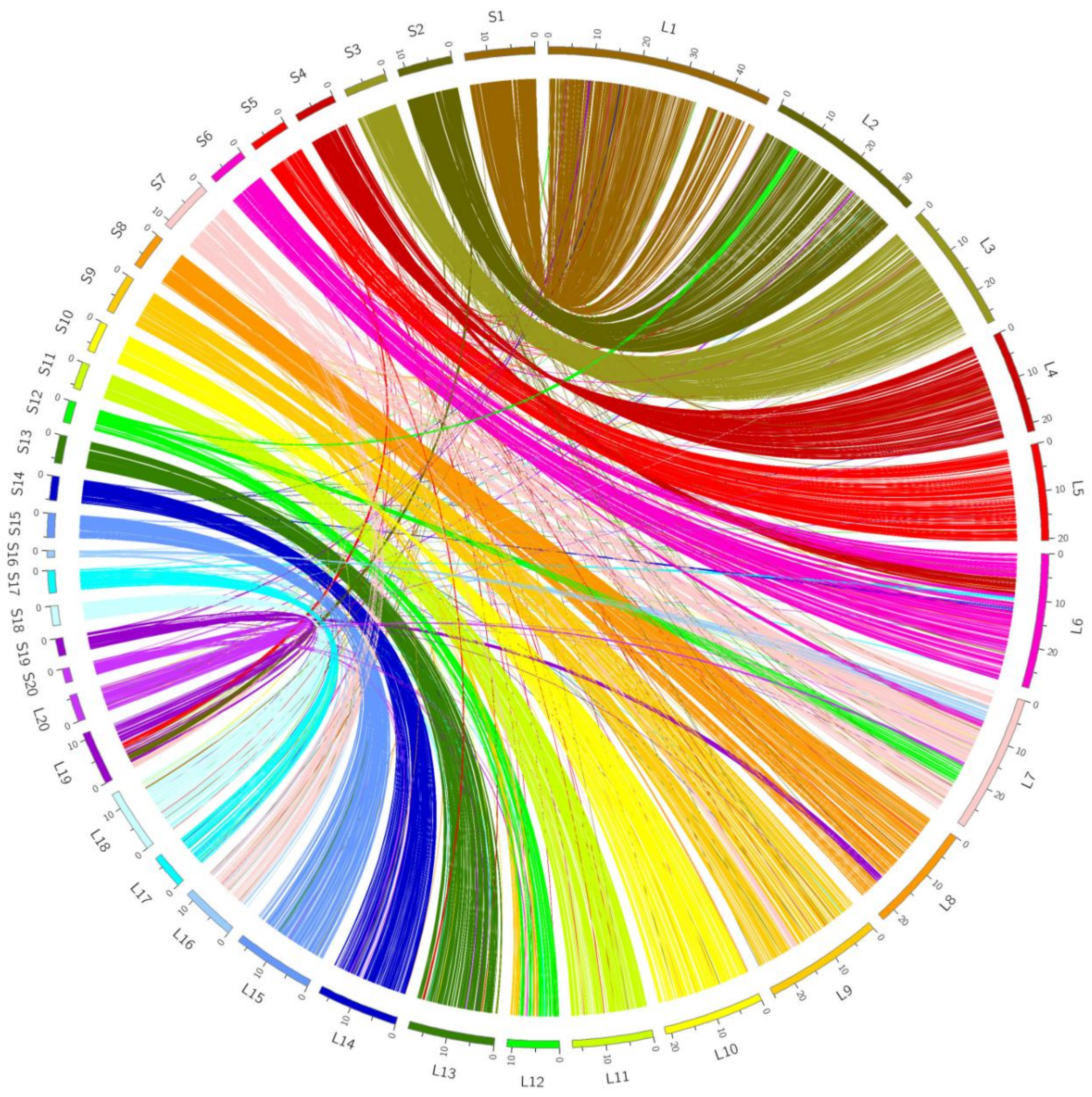


**Supplementary Fig. 5.**

Syntenic circos plot of genomic base-pairs of *Landoltia punctata* strain 0202 and *Spirodela polyrhiza* strain 9509.

The “L” and “S” associating with a number present the chromosome of *Landoltia punctata* and *Spirodela polyrhiza*, respectively.

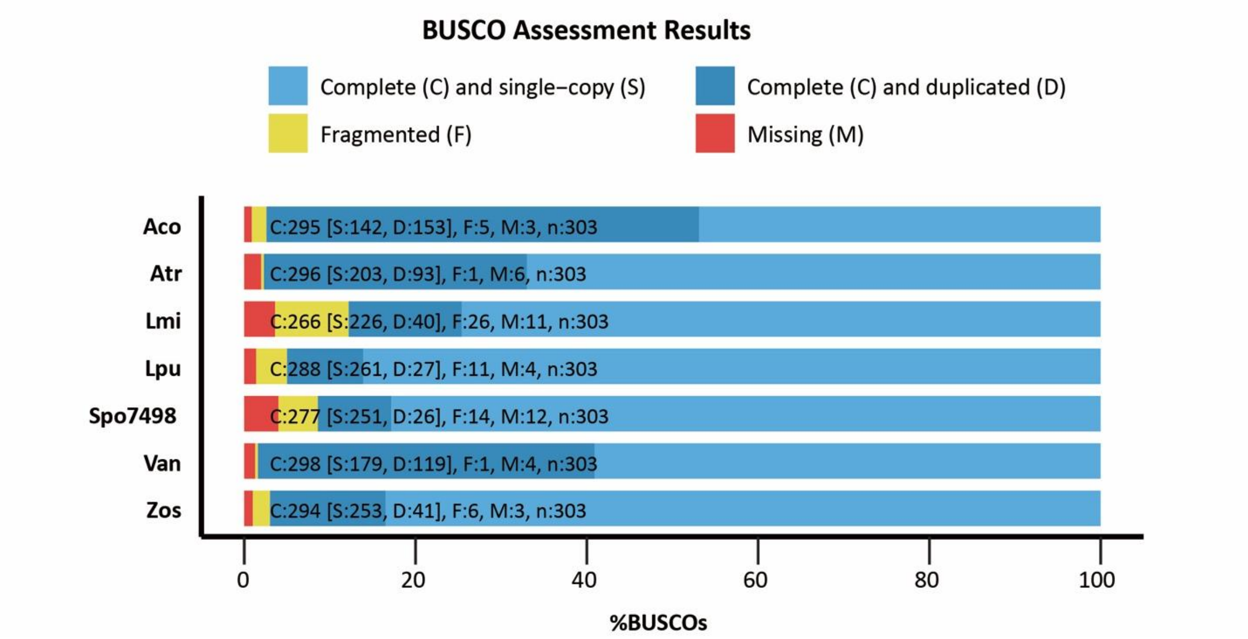


**Supplementary Fig. 6.**

Assessment of genome annotation using BUSCO.

Aco, *Ananas comosus* (L.); Atr, *Amborella trichopoda*; Lmi, *Lemna minor*; Lpu, *Landoltia punctata*; Spo7498, *Spirodela polyrhiza* strain 7498; Van, *Vigna angularis*; Zos, *Zostera marina*.

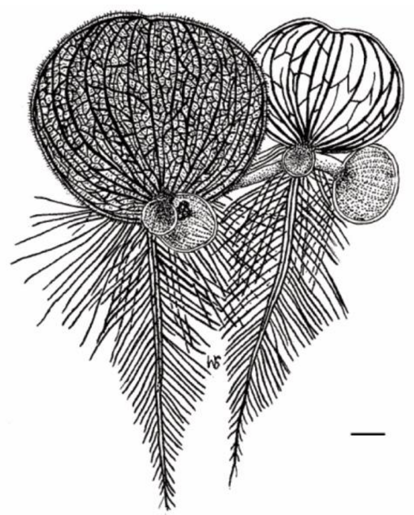

**Supplementary Fig. 7.**

The fossil reconstruction of *Limnobiophyllum scutatum*[1].

Scale bar, 2 cm.

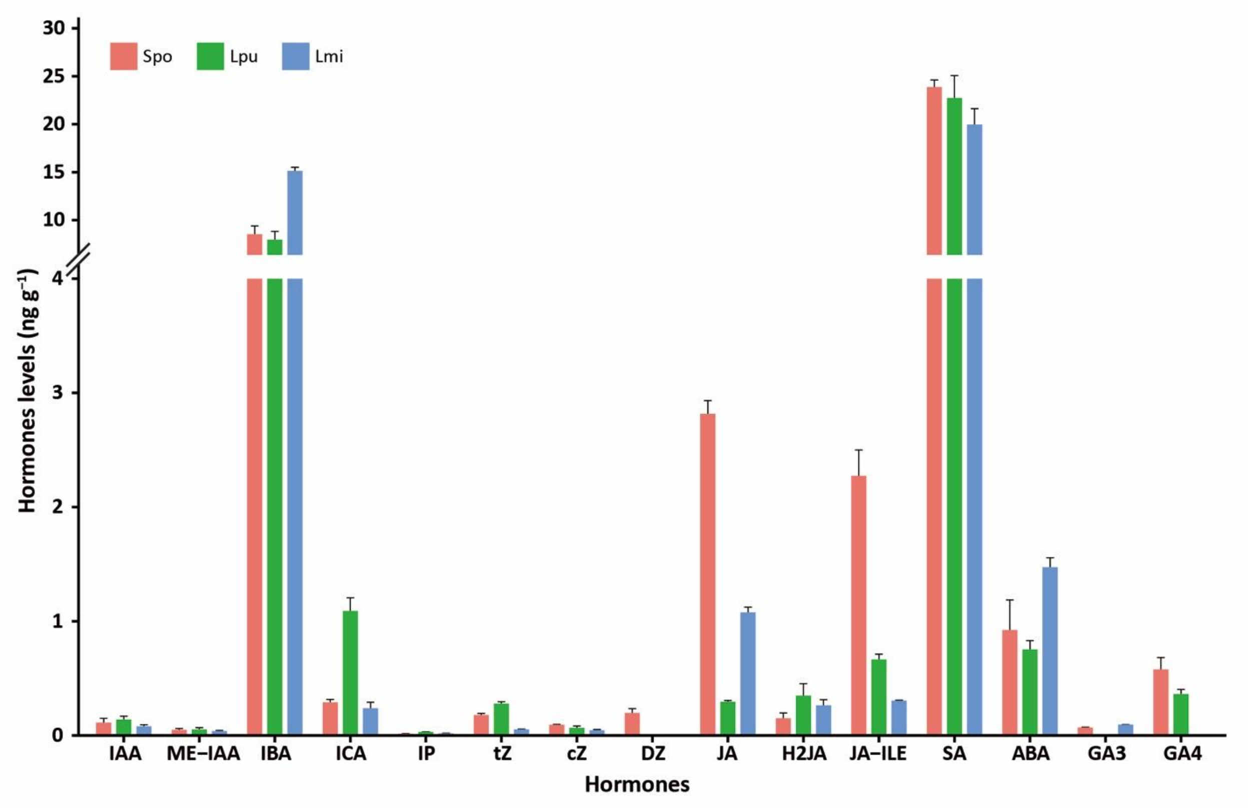


**Supplementary Fig. 8.**

Detection of phytohormones in duckweeds.

Spo, *Spirodela polyrhiza*; Lpu, *Landoltia punctata*; Lmi, *Lemna minor*. IAA, indole-3-acetic acid; ME-IAA, methylindole-3-acetic acid; IBA, indole-3-butyric acid; ICA, indole-3-carboxaldehyde; IP, isopentenyl adenine; tZ, trans-Zeatin; cZ, cis-Zeatin; DZ, dihydro-zeatin; JA, jasmonate acid; H2JA, dihydro jasmonic acid; JA-ILE, jasmonic acid-isoleucine; SA, salicylic acid; ABA, abscisic acid; GA3, gibberellin 3; GA4, gibberellin 4.

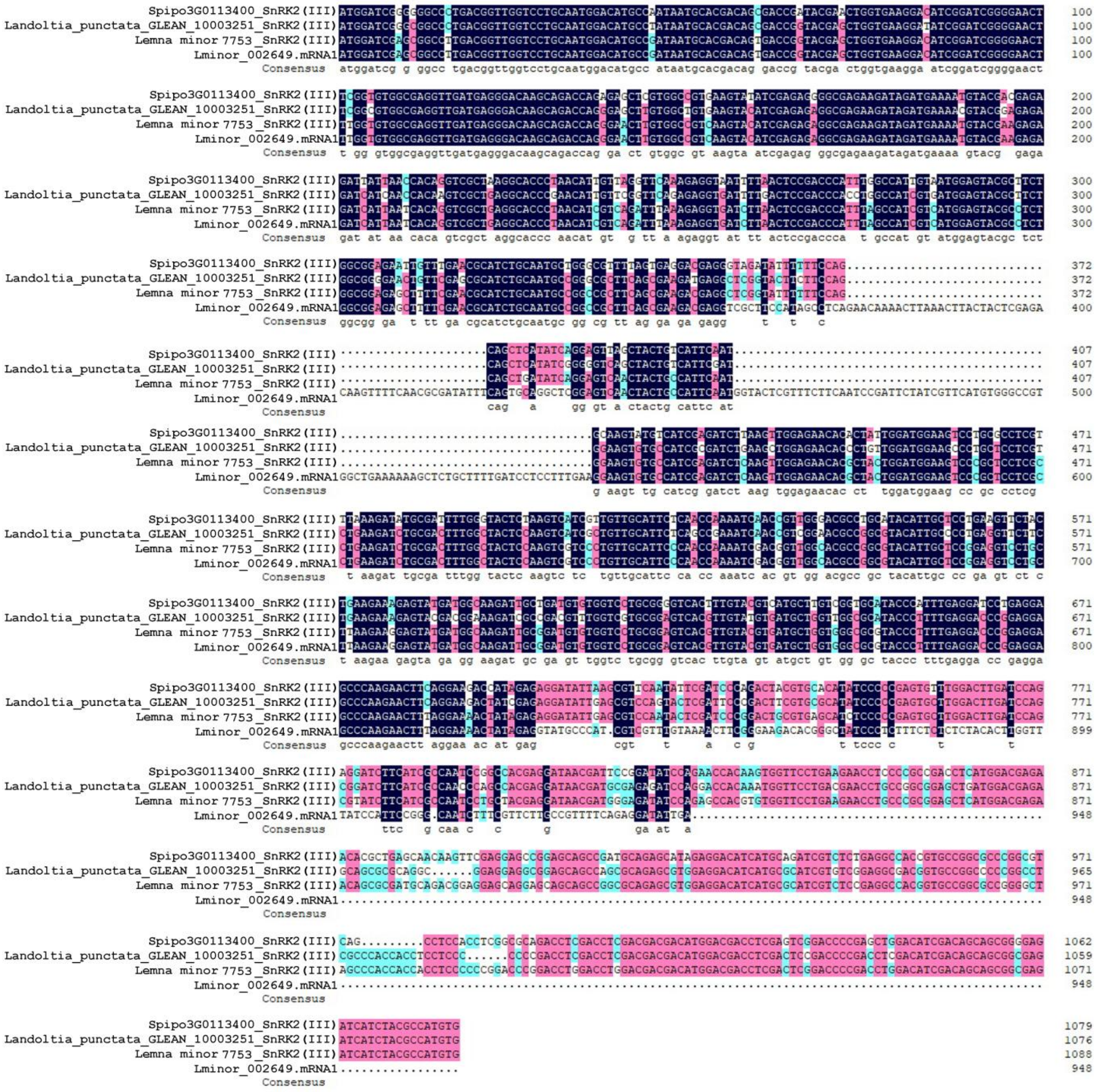

**Supplementary Fig. 9.**

The comparison of SnRK2 (III) coding sequence in *Spirodela polyrhiza*, *Landoltia punctata*, *Lemna minor* and *Lemna minor* reference genome[2].

Spipo3G0113400_SnRK2 (III) is the Group III SnRK2 in *Spirodela polyrhiza*, Landoltia_punctata_GLEAN_10003251_SnRK2 (III) is the Group III SnRk2 in *Landoltia punctata*, Lemna minor 7753_SnRK2 (III) is the Group III SnRK2 in *Lemna minor* and Lminor_002649.mRNA1 is the suspected Group III SnRK2 in *Lemna minor* reference sequence published by[2].

**Supplementary Tables**

**Supplementary Table 1.**

Statistics of raw and filtered genomic sequencing data.

PE, paired-end libraries; MP, mate-pair libraries.

| Library | Insert Size (bp) | Library type | Read Length (bp) | Raw Data (Gb) | Clean data (Gb) | Read Q20 (read1; read2) (%) | Coverage (×) |
|---|---|---|---|---|---|---|---|
| wHAXPI000147-26 | 200 | PE | 100 | 19.0 | 16.2 | 99.8; 99.7 | 80.6 |
| wHAXPI009506-104 | 200 | PE | 100 | 19.8 | 18.4 | 99.7; 99.6 | |
| wHAIPI000148-97 | 500 | PE | 100 | 16.1 | 12.0 | 99.8; 98.5 | 55.6 |
| wHAIPI009496-105 | 500 | PE | 100 | 14.1 | 11.9 | 99.8; 98.0 | |
| wHAMPI012249-33 | 800 | PE | 100 | 13.2 | 11.7 | 99.8; 98.5 | 51.8 |
| wHAMPI012250-33 | 800 | PE | 100 | 13.0 | 10.6 | 99.8; 95.9 | |
| WHLEMdamDEAADWAAPEI-30 | 2000 | MP | 90 | 8.7 | 4.8 | 99.8; 98.6 | 11.1 |
| WHLEMdamDEABDLAAPEI-31 | 5000 | MP | 90 | 8.1 | 3.6 | 99.8; 98.2 | 8.4 |
| WHLEMdamDGAADTAAPEI-11 | 10,000 | MP | 50 | 15.1 | 2.0 | 99.7; 92.7 | 4.6 |
| WHLEMdamDHAADUAAPEI-8 | 20,000 | MP | 50 | 18.0 | 2.6 | 99.9; 97.1 | 12.4 |
| WHLEMdamDHABDUBAPEI-9 | 20,000 | MP | 50 | 21.3 | 2.8 | 99.8; 97.1 | |
| Total | | | | 166.4 | 96.6 | | 224.4 |

**Supplementary Table 2.**

Result of 17-mer frequency distribution analyses.

| Insert size (bp) | K-mer number | Peak depth | Estimated genome size |
|---|---|---|---|
| 200 | 29,071,396,956 | 70 | 415,305,670 |
| 500 | 20,058,033,744 | 48 | 417,875,703 |
| 800 | 18,671,762,844 | 45 | 414,928,063 |

**Supplementary Table 3.**

Summary of assembly statistics of *Landoltia punctata*.

| | Contigs | | Scaffolds | |
|---|---|---|---|---|
| | Length (bp) | Number | Length (bp) | Number |
| L50 | 54,011 | 2,164 | 4,030,252 | 30 |
| Max. Length | 1,111,587 | - | 17,897,791 | - |
| Number (≥100 bp) | - | 63,986 | - | 48,966 |
| Number (≥2000 bp) | - | 12,630 | - | 745 |
| Total size | 403,354,422 | - | 422,361,459 | - |
| GC content | 39.1% | | 36.5% | |

**Supplementary Table 4.**

Comparison of assembly quality between *Landoltia punctata* and other plants with published genome.

-, data not available.

| Species | Contig L50 | Scaffold L50 | Genome size Assembly | Reference |
|---|---|---|---|---|
| *Landoltia punctata* | 54.0 Kb | 4.0 Mb | 422.4 Mb | This Study |
| *Selaginella moellendorffii* | 119.8 Kb | 1.7 Mb | 212.6 Mb | [3] |
| *Musa acuminata* | -[1] | 1.3 Mb | 472.2 Mb | [4] |
| *Oryza sativa L. ssp. indica* | 6.69 Kb | 11.76 Kb | 466 Mb | [5] |
| *Sesamum indicum* L. | 52.2 Kb | 2.1 Mb | 274 Mb | [6] |
| *Phyllostachys heterocycla* | 11.8 Kb | 328 Kb | 2.05 Gb | [7] |
| *Nelumbo nucifera Gaertn* | 38.8 Kb | 3.4 Mb | 804 Mb | [8] |
| *Elaeis guineensis* | - | 1.27 Mb | 1.535 Gb | [9] |
| *Amborella trichopoda* | - | 4.9 Mb | 706 Mb | [10] |
| *Capsicum annuum* | - | 2.47 Mb | 3.06 Gb | [11] |
| *Beta vulgaris* | - | 2.01 Mb | 566.6 Mb | [12] |
| *Spirodela polyrhiza* | - | 3.7 Mb | 145 Mb | [13] |
| *Eucalyptus grandis* | 2.3 Mb | 5 Mb | 605 Mb | [14] |
| *Coffea canephora* | - | 1.26 Mb | 568.6 Mb | [15] |
| *Phalaenopsis equestris* | 20.5 Kb | 359 Kb | 1.086 Gb | [16] |
| *Hordeum vulgare* L. var. nudum | 18.1 Kb | 242 Kb | 3.89 Gb | [17] |
| *Saccharina japonica* | 58.8 Kb | 252 Kb | 537 Mb | [18] |
| *Ananas comosus* (L.) Merr. | 126.5 Kb | 11.8 Mb | 382 Mb | [19] |
| *Lemna minor* | 20.9 Kb | 23.6 Kb | 472Mb | [20] |
| *Oropetium thomaeum* | 2.4 Mb | - | 245 Mb | [21] |
| *Zostera marina* | 79.9 Kb | 485.5 Kb | 202.3 Mb | [22] |

| | | | | |
|---|---|---|---|---|
| *Phaseolus vulgaris* L. | 3.27 Mb | 5 Mb | 587 Mb | [23] |
| *Salvia miltiorrhiza* | 12.38 Kb | 51.02 Kb | 538 Mb | [24] |
| *Hevea brasiliensis* | - | 1.28 Mb | 1.37 Gb | [25] |

**Supplementary Table 5.**

Assessment of genome assembly using expressed sequence tags (ESTs).

| Dataset | Total number | Total Length | Covered by assembly (%) | With >90% sequence in one scaffold | | With >50% sequence in one scaffold | |
|---|---|---|---|---|---|---|---|
| | | | | number | ratio | number | ratio |
| ≥0bp | 3,794 | 2,397,521 | 93.8 | 2,950 | 77.8 | 3,720 | 98.1 |
| ≥200bp | 3,794 | 2,397,521 | 93.8 | 2,950 | 77.8 | 3,720 | 98.1 |
| ≥500bp | 2,862 | 2,040,428 | 94.8 | 2,559 | 89.4 | 2,806 | 98.0 |
| ≥1000bp | 138 | 144,166 | 95.4 | 132 | 95.7 | 133 | 96.4 |

**Supplementary Table 6.**

Summary of assessment of genome assembly with ESTs.

| Total | None | Short | Unique | Multi |
|---|---|---|---|---|
| 3,794 | 21 | 53 | 3,074 | 646 |
| Percent (%) | 0.6 | 1.4 | 81.0 | 17.0 |

**Supplementary Table 7.**

Repetitive sequences annotation in the assembly of *Landoltia punctata*.

| Methods | Repeat size (bp) | Percent (%) |
|---|---|---|
| Homology-based | | |
| TRF | 54,373,159 | 12.9 |
| RepeatMasker | 40,126,737 | 9.5 |
| RepeatProteinMask | 33,969,413 | 8.0 |
| *De novo* | | |
| RepeatModeler | 218,847,027 | 51.8 |
| Total | 250,178,678 | 59.2 |

**Supplementary Table 8.**

Types of transposable elements (TEs) in *Landoltia punctata*.

| Type | Length (bp) | Percent (%) |
|---|---|---|
| DNA transposons | 67,513,324 | 16.0 |
| Retrotransposon | | |
| LINE | 10,390,636 | 2.5 |
| SINE | 162,159 | 0.0 |
| LTR | 81,862,377 | 19.4 |
| Other | 3,698 | 0.0 |
| Unknown | 89,547,421 | 21.2 |
| Total | 221,800,767 | 52.5 |

**Supplementary Table 9.**

Statistics of predicted protein-coding genes in the *Landoltia punctata*.

GN, gene number; AGL, average gene length; ACL, average CDA length; TEx, total exon; AExN, average exon number; AExL, average exon length; AInL, average intron length.

| Gene set | GN | AGL (bp) | ACL (bp) | TEx | AExN | AExL (bp) | AInL (bp) |
|---|---|---|---|---|---|---|---|
| Homolog search | | | | | | | |
| *Arabidopsis. thaliana* | 14,057 | 3,138.2 | 1,140.4 | 72,644 | 5.2 | 220.7 | 479.3 |
| *Oryza. sativa* | 14,830 | 3,118.3 | 1,132.4 | 72,281 | 4.9 | 232.3 | 512.6 |
| *Zea. mays* | 14,334 | 2,807.4 | 1,079.4 | 69,433 | 4.8 | 222.8 | 449.5 |
| *Brachypodium. distachyon* | 13,363 | 3,089.1 | 1,131.7 | 69,075 | 5.2 | 218.9 | 469.5 |
| *Sorghum. bicolor* | 14,625 | 2,888.9 | 1,085.2 | 71,494 | 4.9 | 222.0 | 463.9 |
| *Spirodela. polyrhiza* | 16,174 | 2,923.3 | 1,085.8 | 77,044 | 4.8 | 228.0 | 488.2 |
| *Lemna. minor* | 15,849 | 3,116.4 | 1,135.4 | 79,486 | 5.0 | 226.4 | 493.4 |
| *De novo* | | | | | | | |
| Augustus | 32,896 | 3,051.1 | 968.1 | 129,643 | 3.9 | 245.6 | 708.3 |
| Genscan | 50,098 | 5,503.7 | 1,027.4 | 264,326 | 5.3 | 194.7 | 1,046.8 |
| RNA | | | | | | | |
| EST | 982 | 5,929.6 | 707.4 | 3,764 | 3.8 | 184.6 | 1,843.3 |
| Transcripts | 24,735 | 7,720.2 | 2,640.0 | 217,825 | 8.8 | 299.8 | 650.8 |
| Total gene set | 19,692 | 3,585.9 | 1,142.9 | 99,846 | 5.1 | 225.4 | 600.2 |

**Supplementary Table 10.**

Number of genes with homology or functional classifications by different methods.

| | Annotated number | Percent (%) |
|---|---|---|
| InterPro | 15,176 | 77.1 |
| GO | 11,764 | 59.7 |
| KEGG | 11,633 | 59.1 |
| Swiss-Prot | 11,857 | 60.2 |
| Total annotated | 15,890 | 80.7 |

**Supplementary Table 11. Data links of genomes used herein.**

| Species | Links |
|---|---|
| *Ostreococcus lucimarinus* | ftp://ftp.ncbi.nlm.nih.gov/genomes/all/GCF/000/092/065/GCF_000092065.1_ASM9206v1/GCF_000092065.1_ASM9206v1_genomic.gff.gz |
| *Klebsormidium flaccidum* | http://www.plantmorphogenesis.bio.titech.ac.jp/~algae_genome_project/klebsormidium/kf_download.htm |
| *Chlamydomonas reinhardtii* | ftp://ftp.ncbi.nlm.nih.gov/genomes/all/GCF/000/002/595/GCF_000002595.1_v3.0/GCF_000002595.1_v3.0_genomic.gff.gz |
| *Marchantia polymorpha* | ftp://ftp.ncbi.nlm.nih.gov/genomes/all/GCA/001/641/455/GCA_001641455.1_Mp_v4/GCA_001641455.1_Mp_v4_genomic.gff.gz |
| *Spirodela polyrhiza* | ftp://ftp.ncbi.nlm.nih.gov/genomes/all/GCA/001/981/405/GCA_001981405.1_ASM198140v1/GCA_001981405.1_ASM198140v1_genomic.gff.gz |
| *Landoltia punctata* | https://www.ncbi.nlm.nih.gov/bioproject/PRJNA546087 |
| *Zostera marina* | ftp://ftp.ncbi.nlm.nih.gov/genomes/all/GCA/001/185/155/GCA_001185155.1_Zosma_marina.v.2.1/GCA_001185155.1_Zosma_marina.v.2.1_genomic.gff.gz |
| *Selaginella moellendorffii* | ftp://ftp.ncbi.nlm.nih.gov/genomes/all/GCF/000/143/415/GCF_000143415.3_v1.0/GCF_000143415.3_v1.0_genomic.gff.gz |
| *Lemna minor* | https://genomevolution.org/GenomeInfo.pl?gid=27419 |
| *Nelumbo nucifera Gaertn* | ftp://ftp.ncbi.nlm.nih.gov/genomes/all/GCF/000/365/185/GCF_000365185.1_Chinese_Lotus_1.1/GCF_000365185.1_Chinese_Lotus_1.1_genomic.gff.gz |
| *Amborella trichopoda* | ftp://ftp.ncbi.nlm.nih.gov/genomes/all/GCF/000/471/905/GCF_000471905.2_AMTR1.0/GCF_000471905.2_AMTR1.0_genomic.gff.gz |
| *Arabidopsis thaliana* | ftp://ftp.ncbi.nlm.nih.gov/genomes/all/GCF/000/001/735/GCF_000001735.3_TAIR10/GCF_000001735.3_TAIR10_genomic.gff.gz |
| *Physcomitrella pattens* | ftp://ftp.ncbi.nlm.nih.gov/genomes/all/GCF/000/002/425/GCF_000002425.3_V1.1/GCF_000002425.3_V1.1_genomic.gff.gz |
| *Picea abies* | ftp://plantgenie.org/Data/ConGenIE/Picea_abies/ |

| | |
|---|---|
| *Utricularia gibba* | ftp://ftp.ncbi.nlm.nih.gov/genomes/all/GCA/002/189/035/GCA_002189035.1_U_gibba_v2/GCA_002189035.1_U_gibba_v2_genomic.gff.gz |
| *Phyllostachys heterocycla* | http://202.127.18.221/bamboo/down.php |
| *Eucalyptus grandis* | ftp://ftp.ncbi.nlm.nih.gov/genomes/all/GCF/000/612/305/GCF_000612305.1_Egrandis1_0/GCF_000612305.1_Egrandis1_0_genomic.gff.gz |
| *Oryza sativa* | ftp://ftp.ncbi.nlm.nih.gov/genomes/all/GCF/000/005/425/GCF_000005425.2_Build_4.0/GCF_000005425.2_Build_4.0_genomic.gff.gz |
| *Zea mays ssp. mays* | ftp://ftp.ncbi.nlm.nih.gov/genomes/all/GCF/000/005/005/GCF_000005005.2_B73_RefGen_v4/GCF_000005005.2_B73_RefGen_v4_genomic.gff.gz |
| *Hevea brasiliensis* | ftp://ftp.ncbi.nlm.nih.gov/genomes/all/GCF/001/654/055/GCF_001654055.1_ASM165405v1/GCF_001654055.1_ASM165405v1_genomic.gff.gz |
| *Camellia sinensis* | http://www.plantkingdomgdb.com/tea_tree/data/gff3/ |
| *Populus trichocarpa* | ftp://ftp.ncbi.nlm.nih.gov/genomes/all/GCF/000/002/775/GCF_000002775.3_Poptr2_0/GCF_000002775.3_Poptr2_0_genomic.gff.gz |
| *Apostasia shenzhenica* | ftp://ftp.ncbi.nlm.nih.gov/genomes/all/GCA/002/786/265/GCA_002786265.1_ASM278626v1/GCA_002786265.1_ASM278626v1_genomic.gff.gz |

**Supplementary Table 12. Characteristics of RNA-sequencing data.**

| Species | Reads length (bp) | Raw Data (bp) | Clean Data (bp) | Q20 (%) |
|---|---|---|---|---|
| *Spirodela polyrhiza* | 150 | 28,409,395,500 | 27,070,171,895 | 98.0 |
| *Landoltia punctata* | 150 | 28,529,510,400 | 27,023,351,344 | 98.1 |
| *Lemna minor* | 150 | 26,161,761,000 | 24,960,421,446 | 98.3 |

**Supplementary Table 13. The results of clean reads alignment with rRNA.**

| Species | All Reads Number | Mapped Reads | Unmapped Reads |
|---|---|---|---|
| *Spirodela polyrhiza* | 184,608,218 | 68,173,988 (36.9%) | 116,434,230 (63.1%) |
| *Landoltia punctata* | 186,161,110 | 56,833,736 (30.5%) | 129,327,374 (69.5%) |
| *Lemna minor* | 170,702,602 | 50,181,076 (29.4%) | 120,521,526 (70.6%) |

**Supplementary Table 14. The number of TEs in three duckweed species.**

| Type | Spo (%) | Lpu (%) | Lmi (%) |
|---|---|---|---|
| DNA transposons | NA | 16 | 5.08 |
| Retrotransposon | 13.06 | 21.9 | 31.2 |
| Other | 1.66 | NA | 3.91 |
| Unknown | NA | 21.2 | 21.27 |
| Total | 14.72 | 52.5 | 61.46 |

### Captions for Supplementary Data

Supplementary Data 1. Detailed information and expression level of genes involved in root development.

Supplementary Data 2. Detailed information and expression level of genes involved in stomata development.

Supplementary Data 3. Number of genes involved in phytohormone pathways.

Supplementary Data 4. Number and expression level of genes involved in cellulose biosynthesis.

Supplementary Data 5. Number and expression level of genes involved in hemicellulose biosynthesis.

Supplementary Data 6. Number and expression level of genes involved in lignin biosynthesis.

Supplementary Data 7. Detail information and number of late embryogenesis abundant (LEA) protein genes.

Supplementary Data 8. Total number of transcription factors (TFs) in duckweeds predicted by iTAK.

Supplementary Data 9. Detail information and number of transcription factors (TFs) in bHLH, C2H2 and WRKY family.

Supplementary Data 10. Detail information about GO enrichment analysis of expanded gene families in duckweeds.

Supplementary Data 11. Detail information about KEGG enrichment analysis of expanded gene families in duckweeds.

Supplementary Data 12. The list of gene families enriched in flavonoid, anthocyanin, flavone and flavonol biosynthesis pathways.

Supplementary Data 13. The expanded gene families list of pentatricopeptide repeat (PPR) proteins.

Supplementary Data 14. List of genes involved in the Skp1/Cullin/F-box-type ubiquitin ligase (SCF) core signaling pathway of Auxin, Jasmonic acid and Gibberellic acid.

Supplementary Data 15. Gene number of main components involved in ABA-induced stomatal closure signaling in eight typical species.

Supplementary Data 16. The gene number and gene expression of auxin signaling pathway involved in AR development.

Supplementary Data 17. The number of genes involved in phenylalanine metabolic pathway.

Supplementary Data 18. The content of cellulose, hemicellulose, and lignin of C_reinhardtii, duckweeds and other plants.

Supplementary Data 19. Single copy genes amino acid sequence of eight species used for phylogenetic tree construction.

Supplementary Data 20. The concatenated chloroplast genome sequences of 23 species used for phylogenetic tree construction.
